# Conformational modulation of a mobile loop controls catalysis in the (βα)_8_-barrel enzyme of histidine biosynthesis HisF

**DOI:** 10.1101/2024.06.21.600150

**Authors:** Enrico Hupfeld, Sandra Schlee, Jan Philip Wurm, Chitra Rajendran, Dariia Yehorova, Eva Vos, Dinesh Ravindra Raju, Shina Caroline Lynn Kamerlin, Remco Sprangers, Reinhard Sterner

**Affiliations:** Institute of Biophysics and Physical Biochemistry, Regensburg Center for Biochemistry, University of Regensburg, Universitätsstrasse 31, 93053 Regensburg, Germany; School of Chemistry and Biochemistry, Georgia Institute of Technology, 901 Atlantic Drive NW, Atlanta, GA 30318

**Author notes:** These two authors contributed equally (Enrico Hupfeld, Sandra Schlee).

## Abstract

The overall significance of loop motions for enzymatic activity is generally accepted. However, it has largely remained unclear whether and how such motions can control different steps of catalysis. We have studied this problem on the example of the mobile active site β1α1-loop (loop1) of the (βα)8-barrel enzyme HisF, which is the cyclase subunit of imidazole glycerol phosphate synthase. Loop1 variants containing single mutations of conserved amino acids showed drastically reduced rates for the turnover of the substrates *N*’-[(5’-phosphoribulosyl) formimino]-5-aminoimidazole-4-carboxamide ribonucleotide (PrFAR) and ammonia to the products imidazole glycerol phosphate (ImGP) and 5-aminoimidazole-4-carboxamide-ribotide (AICAR). A comprehensive mechanistic analysis including stopped-flow kinetics, X-ray crystallography, NMR spectroscopy, and molecular dynamics simulations detected three conformations of loop1 (open, detached, closed) whose populations differed between wild-type HisF and functionally affected loop1 variants. Transient stopped-flow kinetic experiments demonstrated that wt-HisF binds PrFAR by an induced-fit mechanism whereas catalytically impaired loop1 variants bind PrFAR by a simple two-state mechanism. Our findings suggest that PrFAR-induced formation of the closed conformation of loop1 brings active site residues in a productive orientation for chemical turnover, which we show to be the rate-limiting step of HisF catalysis. After the cyclase reaction, the closed loop conformation is destabilized, which favors the formation of detached and open conformations and hence facilitates the release of the products ImGP and AICAR. Our data demonstrate how different conformations of active site loops contribute to different catalytic steps, a finding that is presumably of broad relevance for the reaction mechanisms of (βα)8-barrel enzymes and beyond.

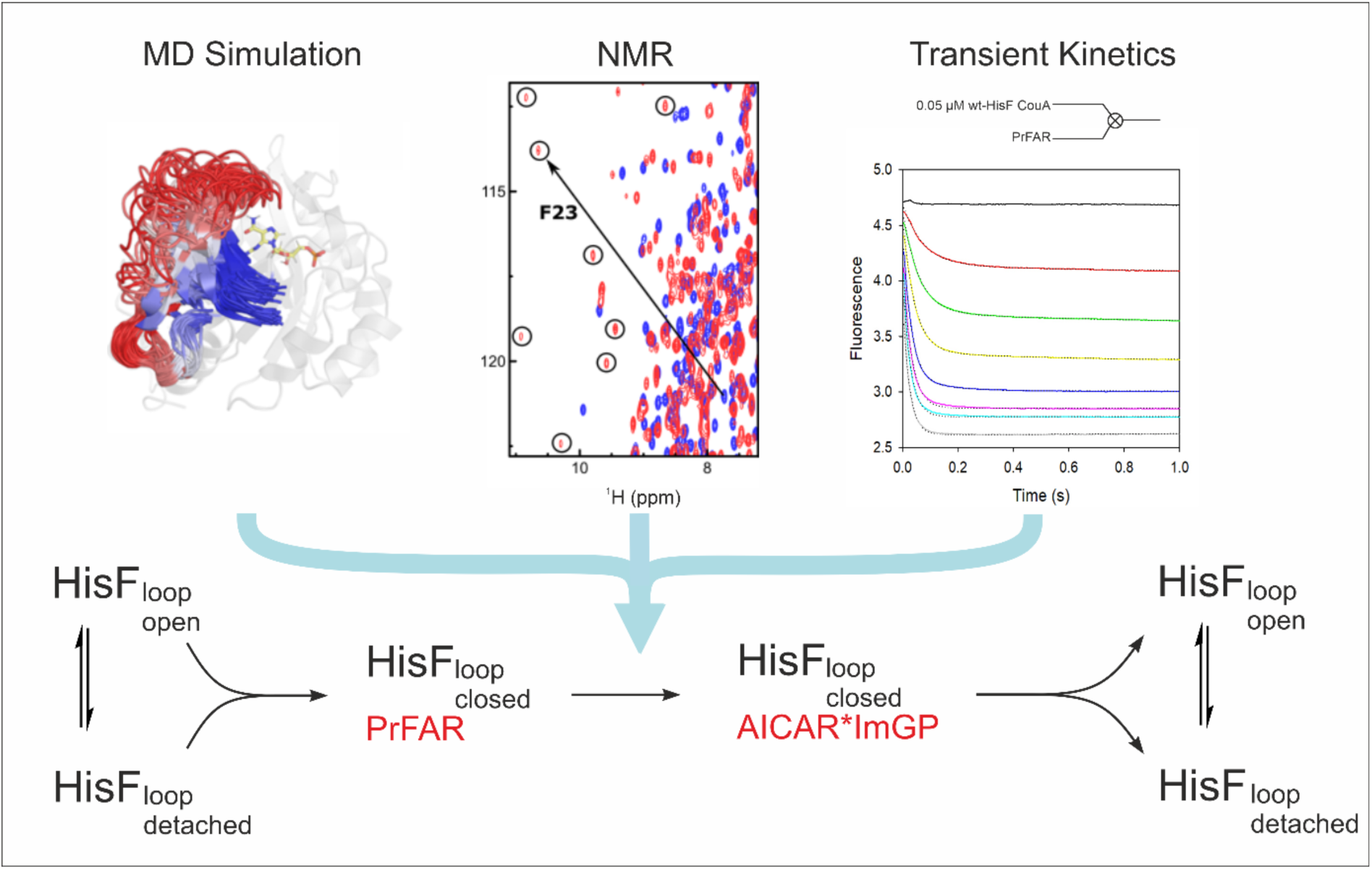

## INTRODUCTION

Enzymes perform reactions with remarkable catalytic efficiency, selectivity and specificity and their function is closely linked to their molecular motions.^1–3^ Along these lines, the process of catalysis typically involves movements of residues in the active site of the enzyme during substrate binding and product release. These steps include motions of single residues as well as opening and closing of loop regions or entire lid domains.^4–5^ Such ligand-driven conformational changes are very well documented^6–7^ and are grouped under the umbrella term “induced-fit motions”, stating that binding of the substrate leads to the transition of non-productive and often poorly defined active site conformations into a single well-defined conformation that is complementary to the reaction transition state.^8–9^ Moreover, it is increasingly recognized that enzyme conformational fluctuations enable the sampling of high-energy intermediates or conformational sub-states along the enzyme reaction coordinate. Experimental evidence continues to indicate that in many cases, the catalytic efficiency of enzymes is directly tied to the rate of conformational transitions into such sub-states.^10–13^ Nevertheless, the direct role of enzyme motions in accelerating the individual states of the catalytic reaction is still under debate.^14–17^

The importance of mobile loops as critical participants in substrate binding and regulation of enzyme activity and specificity is reflected in natural enzyme evolution, where sequence changes are frequently localized at loop regions and the associated modifications of loop conformational plasticity have contributed to diversification of various enzyme families.^18–19^ In addition, it has become increasingly clear that loop mobility needs to be considered in protein engineering approaches aiming at the development of more powerful enzyme catalysts.^20–22^

The (βα)8-or TIM-barrel fold is the most abundant and most versatile fold of enzymes in nature. Around 10 % of all structurally characterized proteins contain at least one domain of this fold.^23^ TIM-barrels catalyse a wide variety of unrelated reactions, covering 5 of the 7 EC classes.^24^ The fold consists of eight alternations of β-strands and α-helices, the strands forming a central β-barrel, which is surrounded by the α-helices. On the C-terminal face of the barrel, the connecting βnαn-loops often contain residues involved in substrate binding and catalysis, while the αnβn+1-loops on the opposite N-terminal face of the barrel mainly contribute to protein stability.^25^ This separation of function and stability is probably one source of the folds versatility and has distinguished it as promising protein scaffold for enzyme design.^26–28^ As the βnαn-loops can be easily modified or exchanged without compromising stability of the protein core, (β/α)8-barrel enzymes are highly suitable for studying the relationship between loop dynamics and catalysis.^29–30^

(βα)8-barrel enzymes, where ligand-induced loop motion has been shown to be a critical component for catalytic activity, include triose phosphate isomerase (TIM) and a number of (βα)8-barrel enzymes involved in tryptophan and histidine biosynthesis, namely TrpF, TrpC, HisA, and PriA.^31–33^ Both the position and length of the loops involved in catalysis, as well as the nature of loop motions and their role in the catalytic mechanism, differ in the various (βα)8-barrel enzymes. To extend the spectrum of loop conformational changes and their relation to the catalytic mechanism in (βα)8-barrel enzymes more broadly, we have focussed on another enzyme of the histidine biosynthetic pathway, the cyclase subunit HisF of imidazole glycerol phosphate synthase (ImGPS) from the hyperthermophilic bacterium *Thermotoga maritima*. HisF catalyzes the conversion of *N*’-[(5’-phosphoribulosyl)formimino]-5-aminoimidazole-4-carboxamide ribonucleotide (PrFAR) and ammonia that is provided by the glutaminase subunit HisH into imidazole glycerol phosphate (ImGP) and 5-aminoimidazole-4-carboxamide-ribotide (AICAR) (**Figure 1A**).^34–37^

**Figure 1.**
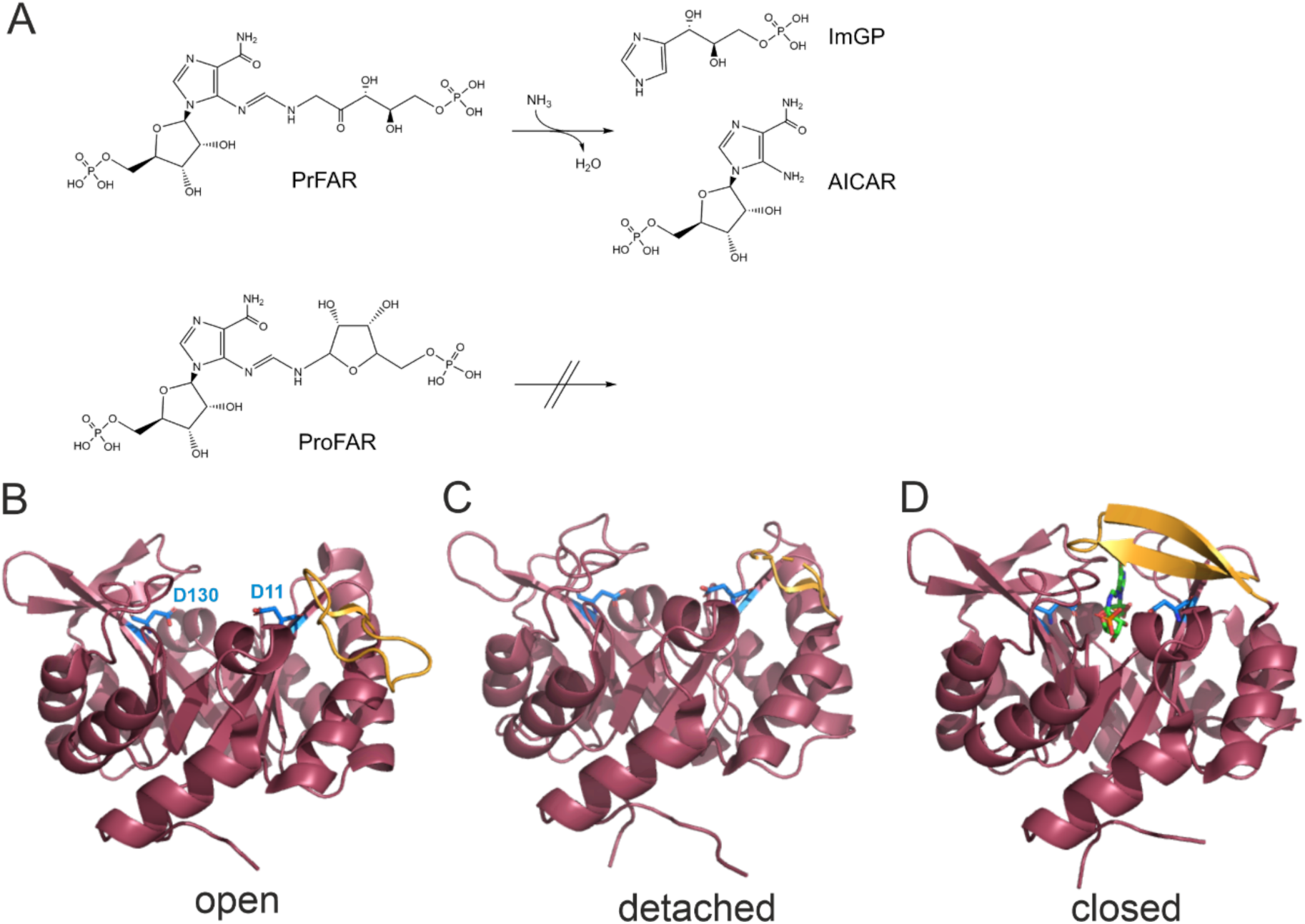
Reaction catalyzed by HisF and crystal structures of HisF from Thermotoga maritima with open/detached/closed loop1 conformations. **(A)** HisF catalyzes the conversion of PrFAR and ammonia into ImGP and AICAR. The substrate analogue ProFAR binds to the active site but is not cleaved. **(B)** In the apo state loop1 (residues R16-G30, orange) adopts an open conformation (PDB entry 1vh738). The catalytic residues D11 (general base, located within β-strand β1) and D130 (general acid, located within β-strand β5) are shown as blue sticks. (C) In some structures of HisF, loop1 is not resolved and presumably detached from the HisF core (PDB entry 3zr439). (C) In the presence of HisH and bound substrate analogue ProFAR (green sticks), loop1 adopts a closed conformation (PDB entry 7ac840, chain E) and forms a β-sheet.

While ImGP is further processed to histidine, the second product AICAR is salvaged in purine biosynthesis. Prokaryotic ImGPS enzymes (including the ImGPS from *Thermotoga maritima)* form heterodimeric bi-enzyme complexes that consist of the cyclase HisF and the glutaminase HisH subunit, which supplies ammonia by glutamine hydrolysis.^41–42^ ImGPS is a well-known model system for studies of allostery.^40, 43–47^ Binding of the substrate PrFAR or its analogue, *N*’-[(5’-phosphoribosyl)formimino]-5-aminoimidazole-4-carboxamide-ribonucleotide (ProFAR)^39^ (**Figure 1A**), results in the allosteric stimulation of the glutaminase reaction, involving a drastically increased rate of glutamine turnover by HisH. Importantly, under *in vitro* conditions the cyclase subunit HisF is able to catalyze the cyclase reaction in the absence of HisH, using externally added ammonium salts at basic pH values.^37^ The active site of HisF is located at the C-terminal face of the central β-barrel, where two conserved aspartate residues, D11 and D130, catalyse the cyclase reaction.^34^ Numerous crystal structures of the isolated subunit HisF^38, 48^ and of the HisH-HisF heterodimer in absence and presence of ligands^39–40, 49^ have been determined. Based on the conformation of the loop that connects strand β1 with helix α1 (loop1), these structures can be separated into an open conformation, where loop1 is flipped toward the outer a-helical barrel ring (**Figure 1B**), a detached conformation, where loop1 is not visible in the electron density and presumably flexible (**Figure 1C**) and a closed conformation, where loop1 closes over the active site (**Figure 1D**).

Here, we aimed at linking the different loop conformations in HisF with function. To that end we combined an extensive mutational analysis with steady-state and stopped-flow enzyme kinetics, NMR spectroscopy, X-ray crystallography, and molecular dynamics simulations. Based on that, we establish a model in which the function of loop1 is to close around the substrate to stabilize it in the active site pocket such that efficient catalysis can take place. Mutants that fail to form a stably closed loop1 conformation are consequently considerably impaired. Further, we demonstrate that mutations alter the distribution of open-detached-closed conformational states between wild-type and mutated HisF variants, in agreement with prior computational work on triosephosphate isomerase^50^, HisF/TrpF/PriA^32^, and protein tyrosine phosphatases^13, 51^. The wide-spread adoption of such conformational fine-tuning of loop dynamics across unrelated enzymes suggests that such evolutionary conformational modulation is a feature not just of (β/α)8-barrel enzymes, but of loopy enzymes more broadly.

## RESULTS

### Substitution of conserved amino acids in loop1 decrease catalytic activity

The influence of loop1 sequence on HisF function was assessed by mutational analysis. First, a multiple sequence alignment (MSA) was compiled which revealed that most residues within loop1 are highly conserved (**Figure 2A**), indicating a function of this loop in the catalytic mechanism. To test this hypothesis, conserved residues were replaced by either alanine, proline, or glycine. Whereas alanine substitutions should uncover effects based on electrostatic or hydrophobic interactions, proline or glycine substitutions were introduced to reveal effects related to loop mobility. The assumption was that introduction of proline residues would render loop1 more rigid in the detached state, whereas inclusion of glycine residues would increase loop1 mobility. The resulting HisF loop1 variants were expressed in *E. coli*, purified, and characterized by steady-state enzyme kinetics. The determined turnover numbers (*k*_cat_) and Michaelis constants for PrFAR (*K*M^PrFAR^) are listed in **Table 1**.

**Figure 2.**
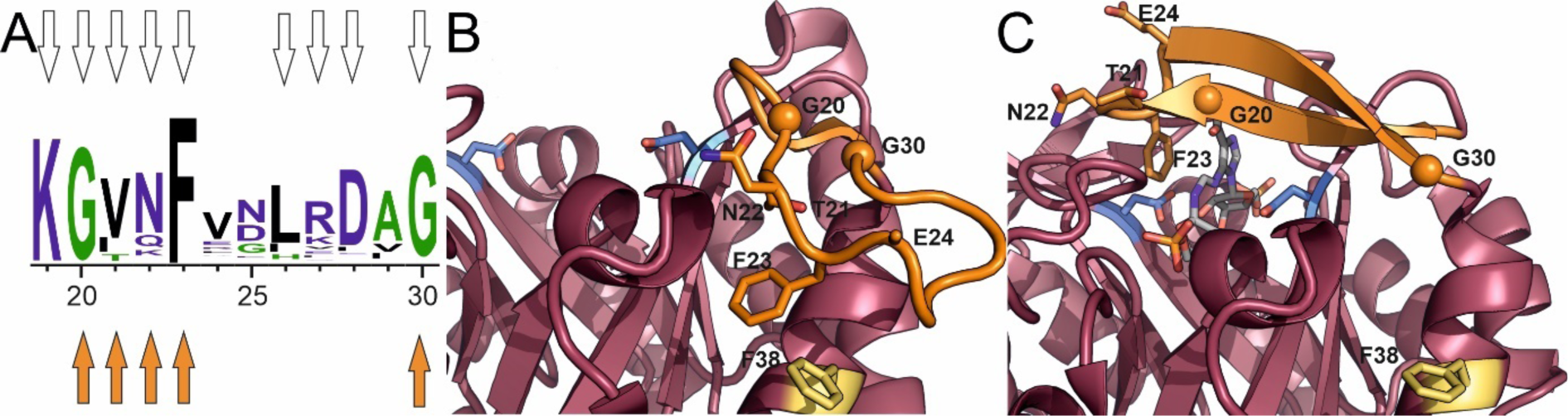
Sequence conservation and mutational analysis of loop1. (**A**) Sequence logo (generated with WebLogo3.6) based on a multiple sequence alignment (MSA) of about 1300 HisF sequences. Residues are numbered according to HisF from *T. maritima*. Mutated residues are marked with white arrows. Residues whose mutation to Ala, Pro, or Gly resulted in a significant reduction of catalytic activity are marked with orange arrows. (**B**) Detailed view of the open loop1 conformation (PDB ID: 1VH7^38^). Functionally important residues within loop1 are shown as orange sticks or spheres. Residue F38 is marked in yellow sticks, the catalytic residues D11 and D130 are shown as blue sticks. (**C**) Detail view of the closed loop1 conformation (PDB ID: 7AC8^40^, chain E). The bound substrate analogue ProFAR is shown in stick representation (colored by element).

**Table 1.** Steady state kinetic parameters of wt-HisF and loop1 variants at 25 °C.

| | $k_{cat}$<br>(s<sup>-1</sup>) | $K_M^{PrFAR}$<br>(μM) | $k_{cat}/K_M^{PrFAR}$<br>(M<sup>-1</sup> s<sup>-1</sup>) |
| --- | --- | --- | --- |
| wt | 2.4 ± 0.2 | 4.5 ± 0.5 | 5.3 × 10<sup>5</sup> |
| K19A | 1.1 ± 0.1 | 6.1 ± 1.8 | 1.8 × 10<sup>5</sup> |
| G20A | 2.2 ± 0.1 × 10<sup>-2</sup> | 2.1 ± 0.2 | 1.0 × 10<sup>4</sup> |
| G20P | n.d. | n.d. | - |
| T21G | 1.8 ± 0.1 × 10<sup>-2</sup> | 5.0 ± 0.8 | 3.6 × 10<sup>3</sup> |
| T21P | n.d. | n.d. | - |
| N22A | 2.9 ± 0.2 × 10<sup>-1</sup> | 8.4 ± 1.7 | 3.4 × 10<sup>4</sup> |
| F23A | 5.5 ± 0.4 × 10<sup>-3</sup> | 6.8 ± 1.3 | 8.1 × 10<sup>2</sup> |
| E24P | n.d. | n.d. | - |
| L26A | 1.1 ± 0.03 | 2.0 ± 0.3 | 5.5 × 10<sup>5</sup> |
| D28A | 2.4 ± 0.1 | 4.5 ± 0.9 | 5.3 × 10<sup>5</sup> |
| G30A | 1.0 ± 0.1 × 10<sup>-1</sup> | 3.7 ± 1.3 | 2.7 × 10<sup>4</sup> |
| G30P | n.d. | n.d. | - |
| F38A | 2.3 ± 0.1 | 3.5 ± 0.6 | 6.6 × 10<sup>5</sup> |
| wt CouA | 1.5 ± 0.1 | 4.1 ± 0.9 | 3.7 × 10<sup>5</sup> |

n. d.: no activity detectable.
Values ± SE for $k_{cat}$ and $K_M^{PrFAR}$ were determined by fitting with equations 1 and 2 of the mean for technical triplicates.

While some importance has been attributed to the residue corresponding to K19 in the homologous yeast enzyme His7^52^, we did not observe significant loss of activity for the K19A variant. Likewise, the amino acid substitutions L26A and D28A did not affect catalytic activity. However, most of the substitutions (G20A, N22A, F23A, G30A, T21G) resulted in a significant decrease of the *k*_cat_ value, whereas the *K*M^PrFAR^ values did not differ by more than two-fold in comparison to wt-HisF. The most severe effects were observed for the proline substitutions (G20P, T21P, E24P and G30P) which caused a drop of catalytic activity below the detection limit. The relatively constant *K*M^PrFAR^ values and the dramatically decreased *k*_cat_ values imply that loop1 does not contribute significantly to the energetics of substrate binding, but rather plays a role for catalysis. As there are no indications that loop1 residues are directly involved in acid-base catalysis, loop1 must play an indirect role in substrate turnover. To obtain insights into this role we have concentrated on three of the identified HisF variants that likely modulate the conformational landscape of loop1. First, the HisF-F23A variant was selected to enhance the flexibility of loop1. This variant will likely destabilize both the closed and open conformations as F23 stacks onto the PrFAR ligand in the closed state and interacts with F38A to form the open state (**Figure 2B, C**). Second, the HisF-G20P variant was selected to restrict conformational flexibility of loop1 in the detached state. At the same time, this variant will destabilize the closed conformation as residue 20 is part of a β-strand in that state (**Figure 2C**). Finally, the HisF-F38A variant was selected. It contains a mutation outside loop1 and is intended to destabilize the open conformation without effecting the closed conformation. Whereas the G20P and F23A substitutions decrease the *k*_cat_ of wt-HisF by several orders of magnitude, the F38A substitution has no effect on the steady-state catalytic parameters (**Table 1**).

### Amino acid substitutions shift the populations of the loop1 conformations

To assess whether the mutations have an influence on the conformation of loop1 we determined the structures of the HisF-F23A and HisF-G20P variants by X-ray crystallography. In the crystal, the HisF-G20P variant was found in the open conformation, similar as the wt-HisF protein (**Figure S1A**). For the HisF-F23A variant the electron density for residues 20-24 in loop 1 was lacking, indicating that the conformation of loop1 shifted from the open towards the detached state (**Figure S1B**).

To complement these static crystal structures, we subjected the wt-HisF, as well as the variants HisF-F23A, HisF-G20P, and HisF-F38A, to a limited proteolysis analysis. This experiment should provide insights into the conformational mobility of the proteins, since protease cleavage rates depend on the accessibility of the respective target^53–54^ and it has been shown previously, that trypsin specifically cleaves HisF after R27 in loop1.^34^ In our experiments we observed that wt-HisF and the variant HisF-F38A are cleaved at similar rates. The HisF-F23A variant, on the other hand, was cleaved faster, whereas the HisF-G20P variant displayed a reduced cleavage rate (**Figure 3A**). These results are in accordance with the static structures that we solved and that suggested a shift towards the mobile detached state for the HisF-F23A variant and a stably formed open conformation for the HisF-G20P variant.

**Figure 3.**
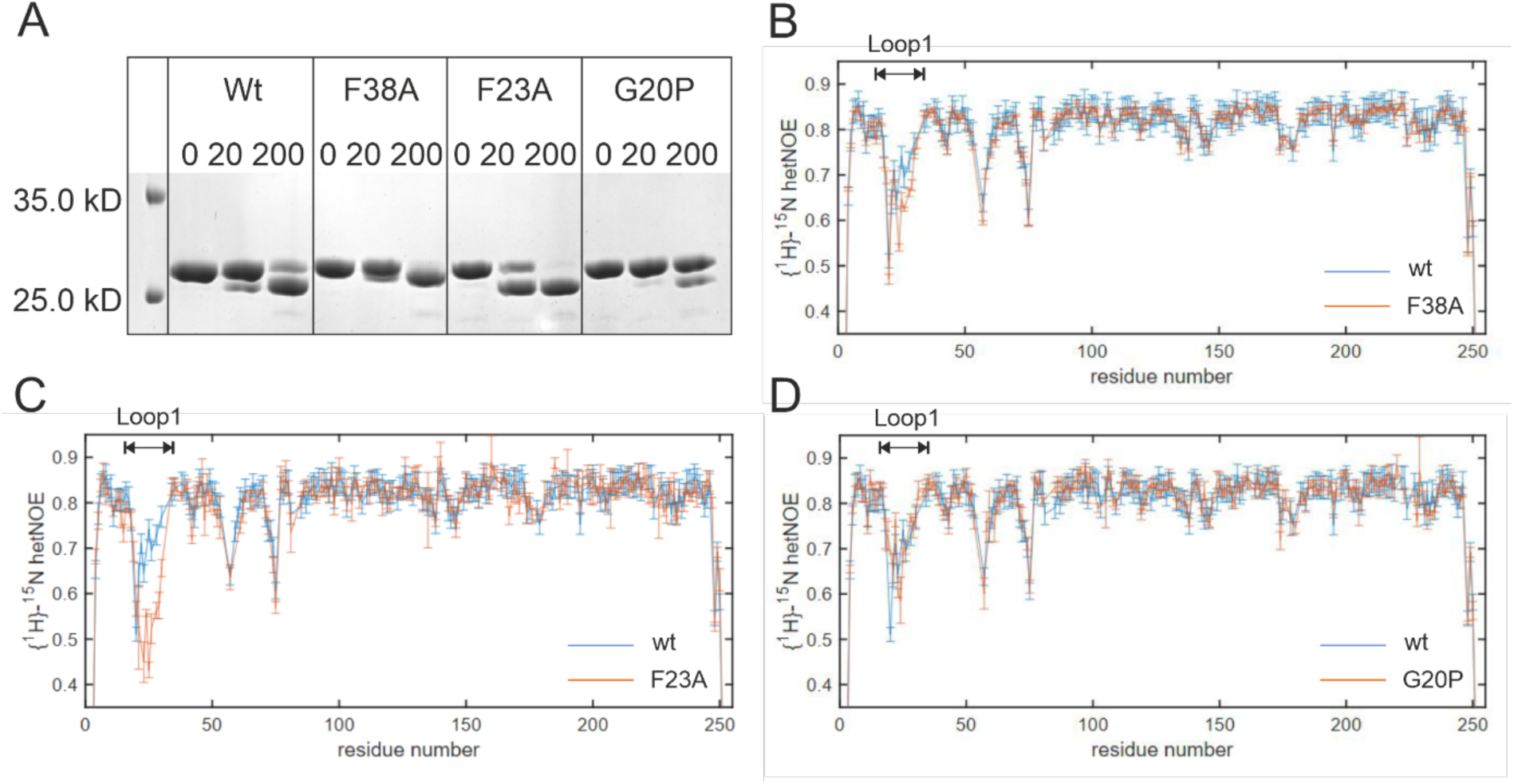
Amino acid substitutions change flexibility and ps-ns dynamics of loop1. **(A)** Limited proteolysis assays monitoring the rates of trypsin cleavage at loop1 residue R27 for wt-HisF, HisF-F38A, HisF-F23A, and His-G20P. Cleavage patterns observed immediately after addition of trypsin (0 min) and after incubation at 25°C for 20 min and 200 min are visualized by SDS polyacrylamide gel electrophoresis (PAGE) analysis. **(B, C, D)** {^1^H}-^15^N hetNOE values of wt-HisF (blue) in comparison to loop1 variants (orange) HisF-F38A **(B)**, HisF-F23A **(C)** and HisF-G20P **(D)**. Decreased values in the loop1 region (residues 19-30) of HisF-F38A and HisF-F23A in comparison to wt-HisF reveal increased dynamics on the ps to ns timescale. The loop dynamics of the HisF-G20P is slightly decreased compared to the wt-HisF.

To obtain direct information on the flexibility of loop1, we exploited NMR experiments. First, we made use of heteronuclear NOE (hetNOE) measurements that probe structural fluctuations on the ps-ns timescale.^55^ These fast motions result in {^1^H}-^15^N hetNOE values below 0.7. For the wt-HisF protein we found that loop1 is the most dynamic loop in the protein. This implies that loop1 predominantly occupies the detached state in solution. It should, however, be noted that the open state of loop1 is also sampled as deletion of loop1 results in chemical shift perturbations in residues that interact with loop1 in the open state. To assess the effect of the mutations on the conformation of loop1 we compared {^1^H}-^15^N hetNOE values of wt-HisF with those of the variants HisF-F38A, HisF-F23A, and HisF-G20P (**Figure 3B-D**). This revealed that the structural flexibility of loop1 is increased in HisF-F23A variant and, to a small degree, in variant HisF-F38A. These findings confirm that loop1 spends more time in the detached state when the open state is destabilized by the F23A and F38A mutations. By contrast, the ps-ns dynamics of loop1 in the HisF-G20P variant is slightly reduced, in agreement with an increased stability of the open state and thus a shift in the conformation away from the detached state.

We further supplemented our analysis with molecular dynamics (MD) simulations of wt-HisF and the HisF-G20P, HisF-F23A and HisF-F38A variants, in both the unliganded state and in complex with PrFAR. In the case of the unliganded enzyme, as there is no experimental evidence for loop closure in this state, we initiated trajectories only from the loop1-open conformation of the enzyme. However, in the case of the PrFAR-bound enzymes, we initiated simulations from both the open and closed states of loop1 for completeness.

**Figure S2** shows the root mean square fluctuations (RMSF) of all Cα-atoms of HisF during MD simulations of the different systems studied. These reflect the flexibility of loop1 (residues 19-30) in wt-HisF and how this is impacted by the mutations. This figure shows only subtle differences in loop1 flexibility among the loop variants: however, given that the loop is highly flexible in all variants, the relative flexibility of the loop will not necessarily change, although there may be shifts within that ensemble between open, detached, and closed states. We further note that the large absolute value of the loop1 RMSF obtained in the simulations of the PrFAR-bound enzymes initiated from the loop1 closed conformation (**Figure S2B**) is due to conformational adjustment of the loop to a new (but still closed) conformation (**Figure S3**), likely due to the change in ligand from ProFAR present in the crystal structure (Figure 1) to the substrate PrFAR (see **Materials and Methods**).

In order to explore the impact of mutations on loop1 flexibility in the different loop states in more detail, we examined the relative mobilities of loop1 based on this RMSF analysis (Figure 4). The mobility data is supplemented by a projection of loop1 motion in wt-HisF along the first principal component, PC1, from principal component analysis (PCA) of these MD simulations to illustrate the dominant dynamic motif. From this data, it can be seen that the relative mobility profiles vary depending on enzyme variant in simulations initiated from the open conformation of loop1 (in both the unliganded and PrFAR bound states of the enzyme, see **Figures 4A and B**), with much more subtle differences in simulations initiated from the PrFAR-bound loop-closed conformation (Figure 4C). This is due to the high mobility of loop1 in all variants, as our simulations shift the loop towards a new closed conformation. We note also that in simulations initiated from the closed state of loop1, we observe a clear monomodal distribution of mobilities in all variants, peaking towards the center of the loop. In contrast, loop mobility is more complex (and variant dependent) in simulations initiated from the loop1 open conformations, likely due to the loop changing shape as it samples both open and detached conformations.

**Figure 4.**
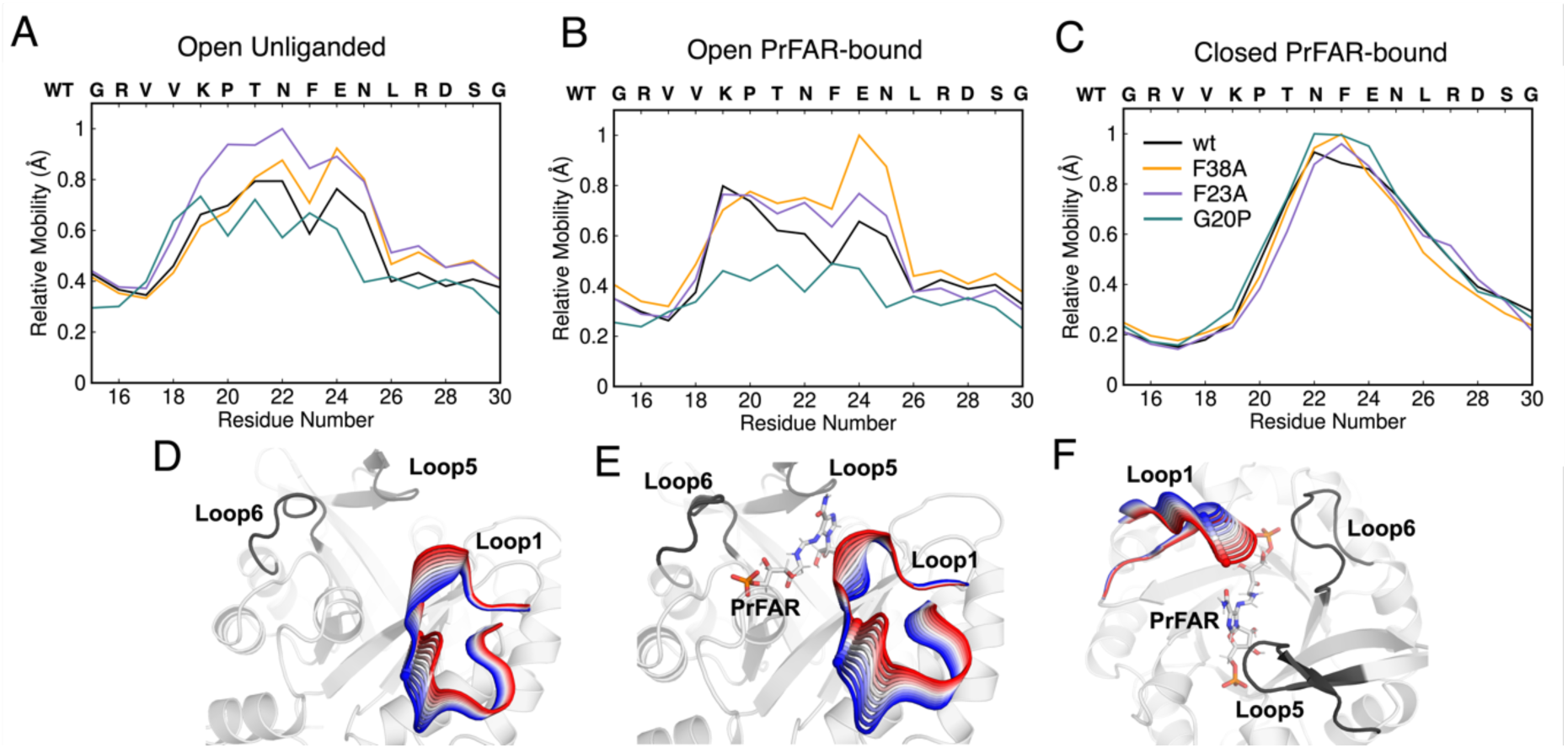
Relative mobility of loop 1 during MD simulations. The mobilities were calculated from the RMSF of the loop1 Cα-atoms, as outlined in the **Supplemental Methods**. Shown here are data from analysis of simulations of the (**A**) unliganded simulations initialized from the loop1 open conformation, and of simulations of the PrFAR-bound enzymes initialized from the loop1 (**B**) open and (**C**) closed conformations. For comparison, panels (**D**-**F**) show projections of the first principal component, PC1, from principal component (PCA) analysis of these simulations (performed as described in the **Supplemental Methods**) onto representative structures of the (**D**) open unliganded, (**E**) open PrFAR bound, and (**F**) closed PrFAR bound states of wt-HisF. The color gradient indicates the transition of loop1 along this principal component.

Finally, to further analyze the flexibility of loop1, we constructed 2D histograms of loop1 motion as a function of the root mean square deviations (RMSD) of the Cα-atoms of loop1 relative to the closed conformation observed in the crystal structure of wild-type HisF/HisH in complex with ProFAR (PDB ID: 7ac8^40^, chain E and F), and the distance RMSD of all non-covalent interactions in the loop1 open conformation of the loop (PDB ID: 1THF^48^) projected as a single vector. The corresponding data is shown in Figures 5, alongside snapshots illustrating the conformational space sampled by loop1 in each set of simulations, colored by Cα-atom RMSF of loop1. From this data, it can be seen that in both unliganded and liganded simulations (Figure 5), we sample both open and detached state (the latter show up as a “smear” on the histograms, as this state is very mobile). The relative population of these states is then shifted by the introduction of point mutations on the loop. In the case of the F38A and F23A variants, we see a clear shift towards more detached states dominating our simulations, but not in the case of the G20P variant. This shift is also illustrated in the enlarged 1D histograms of the dRMSD from the open state contacts (y-axis of the 2D plot), where the histogram of low dRMSD values is decreased for F23A and F38A and increased for G20P. This is in agreement with (and confirming) the observations from our {^1^H}-^15^N hetNOE experiments (Figure 3).

**Figure 5.**
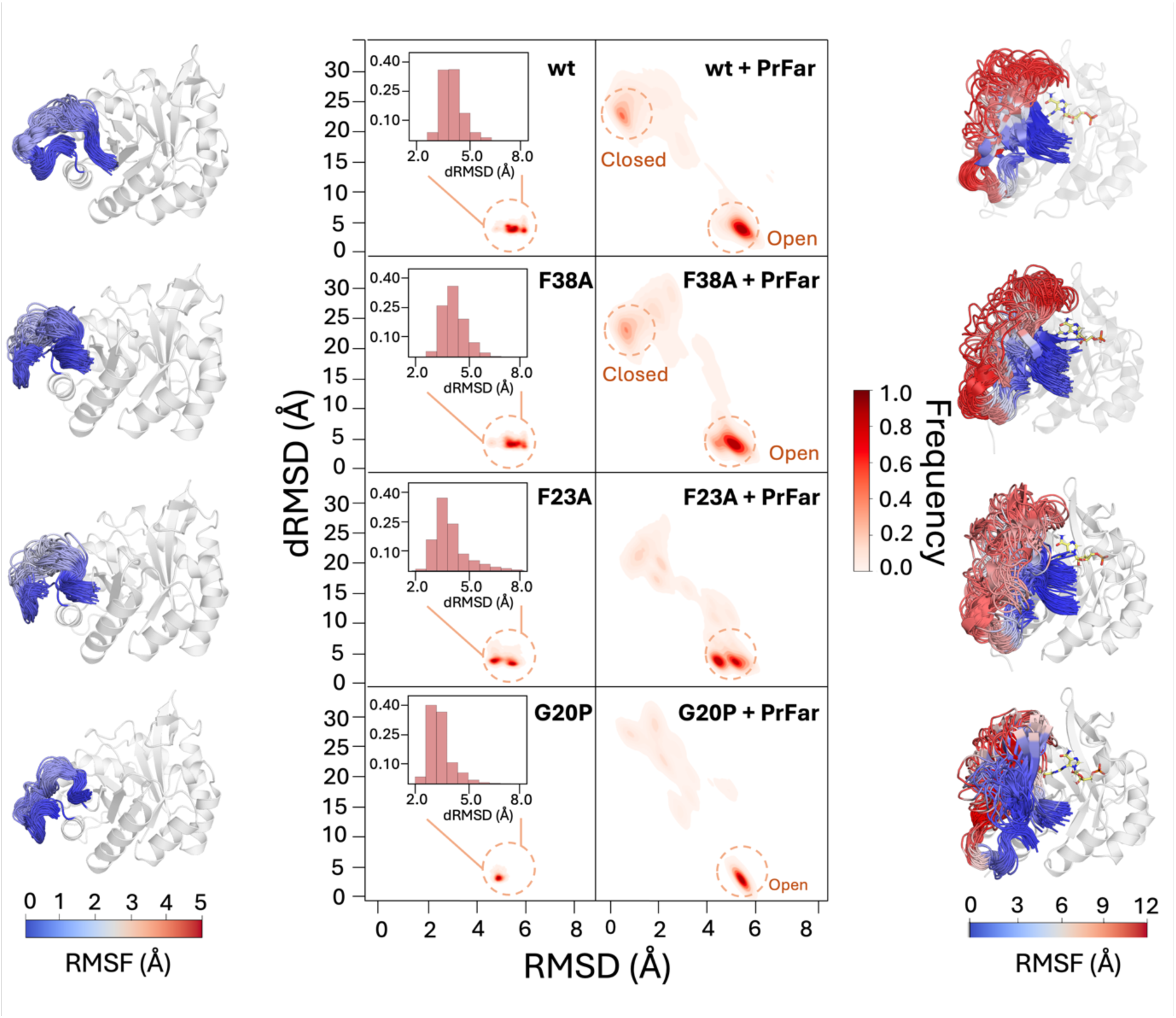
Conformational ensemble of loop1 during molecular dynamics simulations of unliganded and PrFAR-bound wt-HisF, HisF-F38A, HisF-F23A, and HisF-G20P. Shown in the middle of the figure are 2D-histograms of the root mean square deviations (RMSD) of the Cα-atoms of loop1 relative to the crystal closed structure of wild-type and the distance RMSD of all non-covalent interactions in the loop1 that stabilize open liganded conformation during simulations of unliganded (left side) and PrFAR-bound systems (right side). Regions corresponding to closed and open conformations are indicated with a circle. Enlarged 1D histograms of the distance RMSD are included along with the 2D-histograms for the simulations without a ligand. For details of how the distance RMSD values were calculated, see the **Supplemental Methods**. The panel on the left shows, from top to bottom, snapshots of loop1 motion in wt-HisF, HisF-F38A, HisF-F23A, and HisF-G20P during the unliganded simulations, colored by the C_α_-atom RMSF of loop1. The panel on the right shows the analogous data from our corresponding PrFAR-bound MD simulations (variants presented in the same order).

Furthermore, in our simulations of the liganded enzyme (Figure 5), where we also included the closed state of the loop in our simulations, we observe only sparse sampling of this closed state in the F23A and G20P variants compared to the corresponding sampling of the closed state in the F38A variant and the wild-type enzyme. A comparison of Ramachandran plots for glycine and proline in wt-HisF and HisF-G20P (**Figure S4**) shows that angles of G20 sampled in our simulations of wild-type closed or closed active conformations are forbidden by proline Ramachandran plot, explaining why the closed state is so destabilized for the HisF-G20P variant. The impaired sampling of a catalytically competent closed conformation in these variants helps rationalize the diminished/abolished activity observed for these variants in the kinetic data (**Table 1**).

In summary, our crystallography, proteolysis, NMR, and simulation data demonstrate that the HisF-G20P and HisF-F23A variants have opposing effects on the conformation of loop1. In both the HisF-G20P and HisF-F23A variants, there is a shift away from the closed conformations in our simulations, with preferred sampling of detached or open states of the loop. However, whereas loop1 primarily samples the open conformation in the HisF-G20P variant, the loop1 population shifts towards the highly flexible detached conformation in the HisF-F23A variant. As these variants both strongly reduce catalytic turnover (**Table 1**), it is not possible to link the population of open and detached conformations of loop 1 with the rate-limiting step (*k*_cat_) in the turnover reaction. Instead, the G20P and F23A substitutions likely influence turnover *via* alterations in the closed conformation of loop1.

### Amino acid substitutions in loop1 have a limited effect on substrate binding affinities

Since different loop1 conformations in the apo state cannot explain the higher catalytic activities of wt-HisF and HisF-F38A compared to HisF-G20P and HisF-F23A, it was next analyzed whether these amino acid substitutions have consequences for substrate or product binding. To study the thermodynamics and kinetics of PrFAR binding to HisF, fluorescence equilibrium titrations and transient fluorescence kinetic measurements were performed. Although intrinsic fluorescence of the single tryptophan residue 156 of HisF has previously been used as spectroscopic signal transmitter in ligand binding studies^45^, it proved unsuitable for kinetic measurements because of the unspecific fluorescence quenching upon addition of the substrate PrFAR. We sought to avoid this effect by the introduction of an alternative fluorescent probe. The unnatural amino acid L-(7-hydroxycoumarin-4-yl)ethylglycine (CouA) was applied because this probe is relatively small, has good spectroscopic properties and can easily be introduced by genetic code extension.^56–57^ CouA has been used extensively as protein-based fluorescent sensor that reports on protein-ligand interactions^56, 58–60^ and enzyme-substrate binding.^61–62^ As the 7-hydroxycoumarin moiety can exist in a number of tautomeric forms in the ground state, absorption/emission maxima are strongly influenced by environmental factors such as dielectric constant, hydration and pH.^63^ For our purposes CouA was incorporated into HisF in place of a lysine at position 132, a position that is not conserved in HisF sequences and has a distance of ∼ 15 Å to the ligand binding site (Figure 6A). HisF-K132CouA variants (wt, F38A, F23A, G20P) were purified with reasonable yields and the incorporation of CouA was verified spectroscopically (**Figure S5**). Steady-state kinetic parameters of CouA-labeled HisF were virtually identical to those of its non-labelled counterpart (**Table 1**), corroborating that enzymatic activity is not affected by incorporation of the fluorophore. Ligand binding is associated with a decrease of CouA fluorescence emission. Equilibrium titrations with the substrate PrFAR were done in the absence of ammonia to allow for observation of the binding separately from the turnover reaction (Figure 6B).

**Figure 6.**
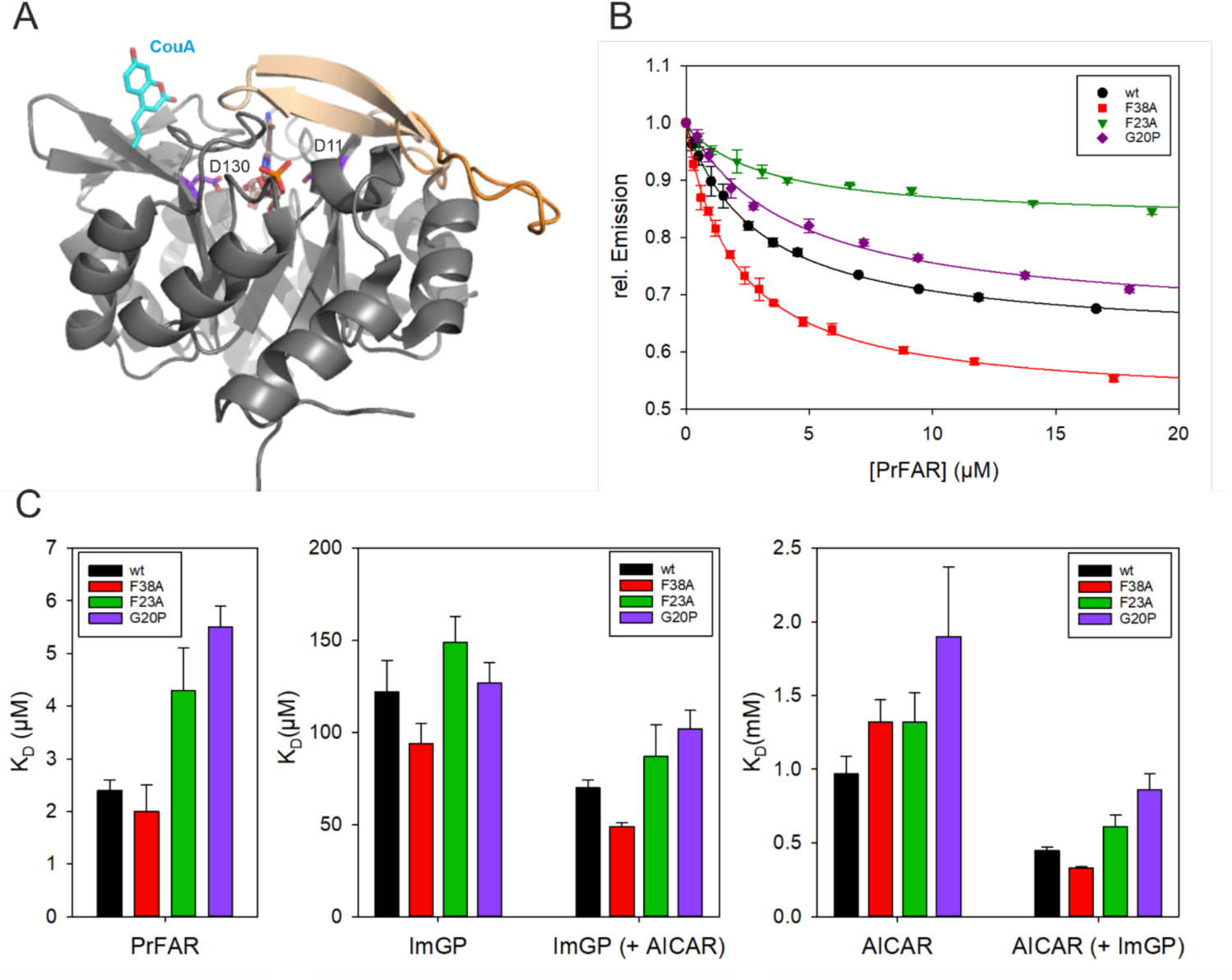
Ligand binding monitored by equilibrium titrations with CouA-labelled HisF. **(A)** Site of CouA incorporation. The structure of HisF is shown with the open loop1 conformation (orange, PDB entry 1vh7^38^) and an overlay of the closed loop1 conformation (beige, PDB entry 7ac8^40^). CouA was modelled into the structure and is shown at position 132 (within β-strand 5) as cyan sticks, the bound substrate precursor ProFAR and the catalytic residues D11 (within β-strand 1) and D130 (within β-strand 5) are shown as sticks. **(B)** Equilibrium titrations of HisF-CouA variants at 25°C. Binding of the substrate PrFAR to the variants (0.2 µM) resulted in a decrease in CouA fluorescence (λ_ex_ = 370 nm, λ_em_ = 452 nm). Relative emission intensity was plotted vs. PrFAR concentration. Lines represent hyperbolic fits of the data. **(C)** Apparent KD values obtained in equilibrium titrations for the binding of the substrate PrFAR or the product molecules ImGP and AICAR to the HisF-CouA variants in absence or presence of the second ligand (AICAR or ImGP), respectively. The associated numerical values are listed in **Table S1**. *K*_D_ values ± SE were determined by fitting the mean ± SEM for at least two technical replicates with equation 3.

The *K*_D_-values determined in the equilibrium titrations show that the G20P and F23A substitutions in loop1 slightly weaken the affinity between HisF and the substrate PrFAR or the products ImGP or AICAR (Figure 6C**, Table S1**). For example, compared between wt-HisF and HisF-F23A the dissociation constant for the substrate PrFAR is increased 1.8-fold, for ImGP 1.2-fold, and for AICAR 1.4-fold. Furthermore, we noticed that the apparent affinity of AICAR is slightly increased in the presence of the second ligand ImGP and vice versa, which indicates a higher formation propensity of the ternary complex (HisF*ImGP*AICAR) in comparison to the respective binary complexes (HisF*AICAR) and (HisF*ImGP). The stabilization effect due to formation of the ternary complex is similar for wt-HisF and HisF-G20P, HisF-F23A, and HisF-F38A, indicating that this is a general feature. Looking at the fluorescence changes upon titration of the dimeric or ternary complexes, another difference between wt-HisF/His-F38A and HisF-F23A/HisF-G20P is noticeable. While the fluorescence amplitudes in the case of HisF-wt and HisF-F38A are higher when the ternary complex is formed than when the binary complexes are formed, the opposite is true for the variants HisF-F23A and HisF-G20P (**Table S2**). This is a first indication that the environment of the fluorophore CouA in the ternary complex for the active variants wt-HisF and HisF-F38A differs from the environment in the inactive variants HisF-F23A and HisF-G20P.

### An induced-fit movement during PrFAR binding is exclusively observed for wt-HisF and HisF-F38A

The kinetics of the PrFAR binding reaction were studied in stopped-flow experiments, whereby the CouA fluorescence decrease was recorded after rapidly mixing the respective HisF-CouA variant with a molar excess of PrFAR. For an assessment of ligand binding kinetics, the observed binding transients were fitted with exponential functions. The number of exponential functions required to describe the transients allows conclusions to be drawn about the number of reaction steps in the binding reaction. In addition, the secondary plots derived from exponential fitting, *e.g. k*obs as function of the PrFAR concentration, provide initial clues to the binding mechanism. In general, time traces (**Figure S6**) for wt-HisF and HisF-F38A resemble each other, whereas time traces associated with the loop1 variants HisF-F23A and HisF-G20P showed notable differences. In the case of wt-HisF (**Figure S6A**) and variant HisF-F38A (**Figure S6B**) time traces are biphasic (sum of two exponential terms). A fast fluorescence decrease is followed by a slow phase with a very small signal amplitude. In the case of loop variants HisF-F23A (**Figure S6C**) and HisF-G20P (**Figure S6D**), single exponential functions were adequate to describe the time traces. The overall fluorescence changes associated with PrFAR binding were smaller, which resulted in lower signal-to-noise ratios. A plot of the first-order rates (*k*_obs_) for the binding reaction as a function of PrFAR concentration provides insights into potential differences in the binding mechanisms of the different variants. In the case of wt-HisF and HisF-F38A, the turnover rate (*k*_obs1_) depends on the substrate concentration in a hyperbolic manner (**Figure S6E**), indicating that binding takes place *via* an induced fit or conformational selection mechanism.^64–65^ In contrast, in the case of loop1 variants HisF-F23A and HisF-G20P, *k*_obs_, increased linearly with increasing concentrations of PrFAR (**Figure S6F**), which is indicative of a simple binding process without involvement of conformational changes. This finding is in accordance with a model where the wt-HisF and HisF-F38A proteins bind the ligand when the protein is in the open or detached conformation, after which loop1 stably closes over the ligand to form the closed conformation. The HisF-G20P and HisF-F23A variants on the other hand are unable to form a stably closed conformation and prefer to remain in the open or detached conformation even in the presence of the ligand, as also observed in our simulations (Figures 5).

To directly assess if the formation of the closed state is impaired in the HisF-G20P and HisF-F23A variants we again turned to NMR titration experiments. To that end, we added ProFAR (a stable PrFAR analogue; see Figure 1A), to ^15^N labelled HisF and followed the induced chemical shift perturbations (CSPs). For all HisF proteins we observed CSPs that directly report on the interactions between HisF and the ligand. Interestingly, we observed a new set of signals that likely reports on the closed conformation of loop 1, as F23 is one of the residues that displays a novel conformation upon PrFAR binding (Figure 7, **circles**). This new set of signals thus reports on the formation of the closed state of loop1 in the presence of the ligand analogue. This stable set of signals does not appear in the HisF-G20P and HisF-F23A variants, proving that the closed conformation is not stably adopted in those cases. This agrees well with simulation data presented in Figure 5.

**Figure 7.**
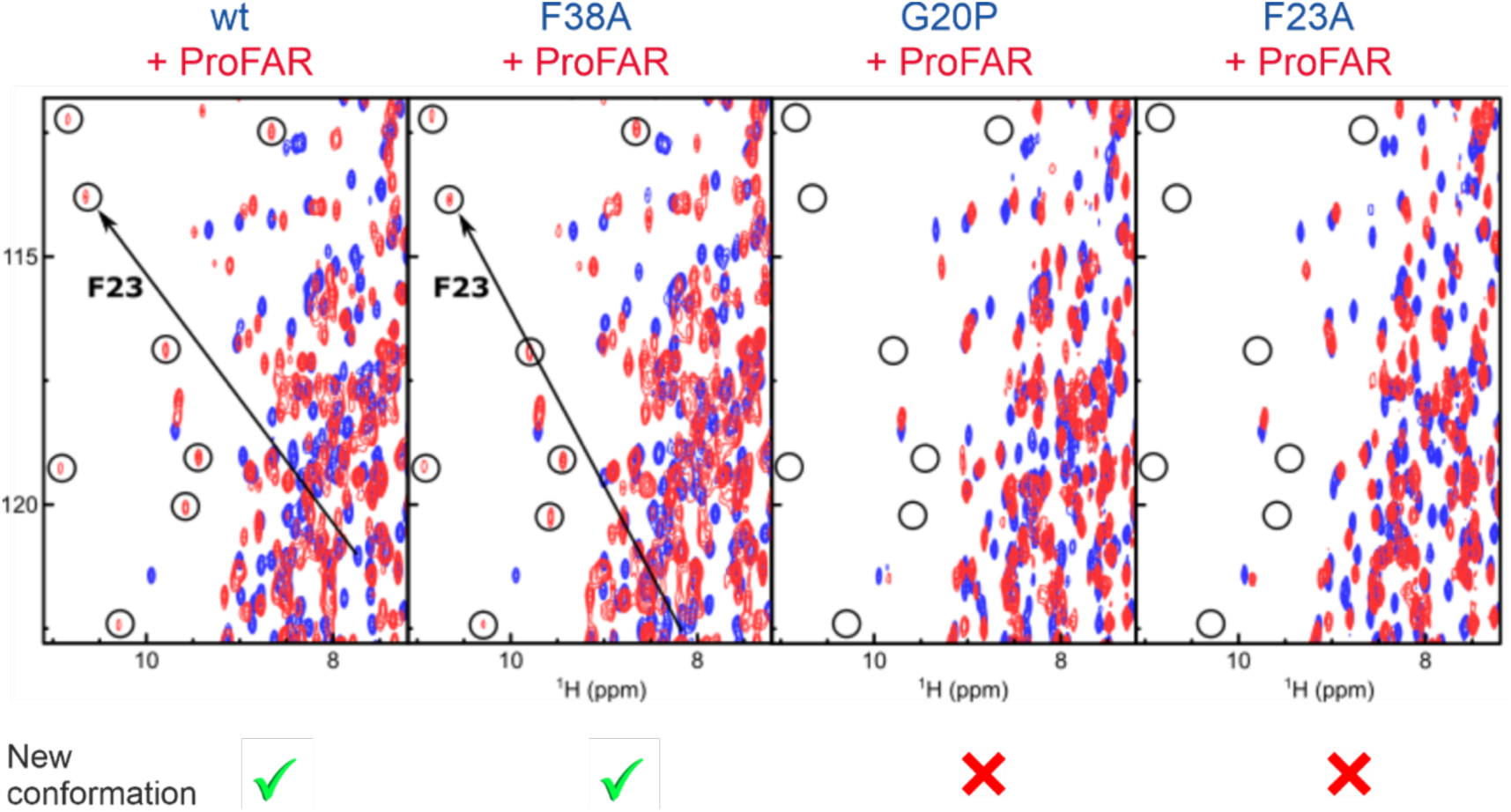
ProFAR binding induces a conformational change of loop1 only in wt-HisF and HisF-F38A. NMR titration experiments recorded in ^1^H-^15^N TROSY spectra showing apo HisF (blue) and HisF in the presence of saturating amounts of ProFAR (red). The large CSP of F23 upon ProFAR binding is shown by a black arrow. The position of several signals with large CSPs is indicated by black circles. Large chemical shift perturbations associated with a substantial conformational change are only observed in the spectra of wt-HisF and HisF-F38A.

Our combined NMR and stopped flow measurements thus are indicative of a two-step binding mechanism in the wt-HisF protein, in which the ligand first interacts with the open or detached conformations of HisF after which loop1 closes to facilitate catalysis (**Scheme 1, top**).

**Scheme 1.**
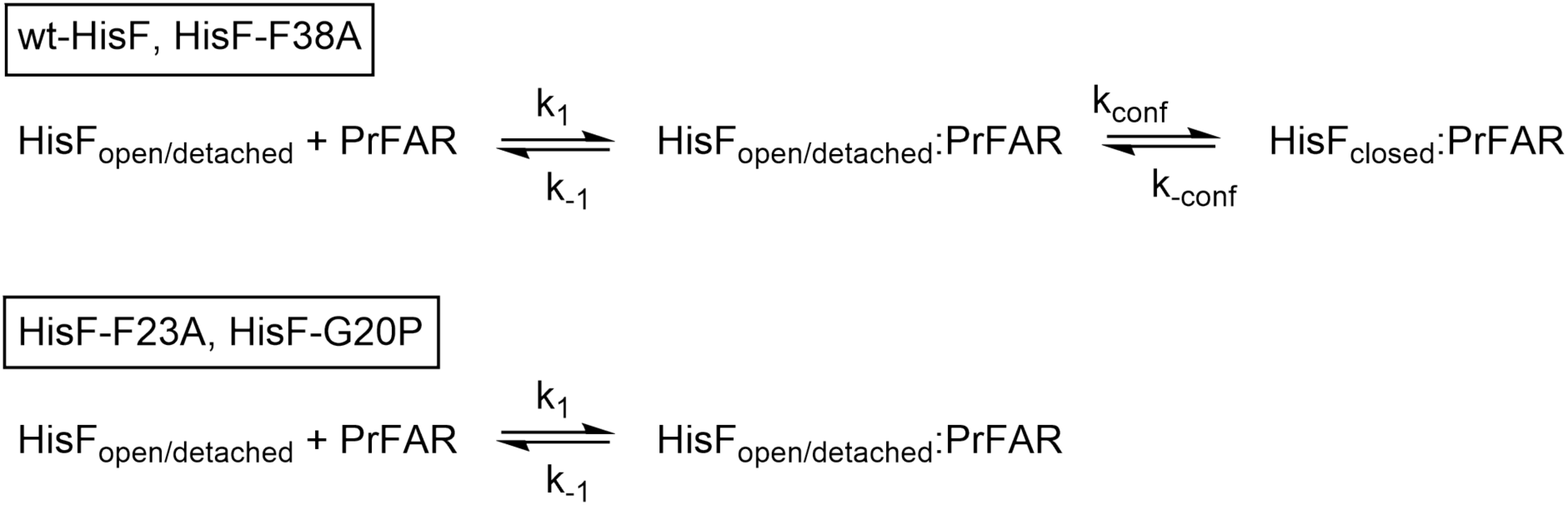
**Binding of PrFAR to wt-HisF and loop1 variants: Induced fit model versus two-state model**

To obtain insights into the rates that are associated with this two-step binding process we fitted the stopped-flow experiments that were performed under pseudo-first order conditions for HisF-CouA (excess of HisF-CouA over PrFAR) to the induced-fit model in **Scheme 1**. These hyperbolic fits allowed for the determination of the *K*_D1_ (= *k*-1/*k*1), *k*_conf_ and *k*_-conf_ for wt-HisF and HisF-F38A (**Table S3**). This shows that the equilibrium of the conformational change is on the closed side and that a stable closed conformation is thus efficiently formed which subsequently facilitates efficient substrate turnover. In contrast, PrFAR binding kinetics for the loop1 variants HisF-F23A and HisF-G20P are compatible with a simple one-step binding reaction (**Scheme 1, bottom**). The *k*_1_ and *k*_-1_ values shown in **Table S3** result from the slope and intercept with the y-axis of a linear fit (**Figure S6F**).

In summary, the NMR and stopped flow experiments, as well as molecular dynamics simulations, establish that substrate binding to HisF occurs via an induced fit mechanism for wt-HisF and for the HisF-F38A variant. The HisF-G20P and HisF-F23A variants on the other hand interact with the substrate *via* a one step binding mechanism as loop1 is, in those cases, unable to close properly over the substrate. Interestingly, these variants still interact efficiently with PrFAR, indicating that loop 1 does not contribute considerably to the binding energy of the substrate.

### Product release from wt-HisF and HisF-F38A is more complex than for HisF-F23A and HisF-G20P

Next, we aimed to obtain insights into the release of the AICAR and ImGP products. Equilibrium titration measurements showed that both ligands, AICAR and ImGP, can bind independently to wt-HisF and all loop1 variants, causing a decrease of CouA fluorescence (Figure 6). Exclusively for wt-HisF and HisF-F38A, we noted a significantly higher fluorescence change when the ternary complex was formed than when the binary complexes were formed (**Table S2**), combined with an increase in apparent binding affinity upon formation of the ternary complex (**Table S1**). This suggests that the interaction of either product (ImGP or AIRCAR) does not result in a conformational change in the enzyme, whereas the interaction with both products at the same time does result in the closing of loop1.

To obtain the rates that are associated with product release from wt-HisF and the loop1 variants we made use of stopped-flow measurements. Representative time traces for wt-HisF are shown in **Figure S7A**, the time traces for HisF-F38A resemble those of wt-HisF (data not shown). As the transient kinetic measurements show, formation of the binary HisF*AICAR and HisF*ImGP complexes is completed within the dead time of the stopped flow device. Based on the used enzyme and substrate concentrations and an instrument dead-time of ∼2.0 ms, an observed rate constant *k*_obs_ of greater than 1000 s^-1^ is required to obscure all evidence of association, suggesting that association rate constants for binary complex formation must be ≥ 10^6^ M^-1^ s^-1^. In contrast, when monitoring the formation of the ternary complex, fluorescence changes with rate constants *k*_obs_ in the range of 50 s^-1^ were observed. This is visible from the exponential fluorescence decrease in the stopped-flow transients when the free enzyme interacts with a mixture of both ligands or when the preformed binary complexes are mixed with the second ligand (**Figure S7A**: HisF + ImGP/AICAR, HisF*ImGP + AICAR, HisF*AICAR + ImGP). These data thus agree with the equilibrium titrations (Figure 6) that revealed a synergistic effect when AICAR plus ImGP bind to the enzyme and with the notion that the interaction with both ligands is associated with a conformational change in the enzyme. In contrast, in the case of the loop variants F23A and G20P, both the binary and ternary complexes were formed within the instrument dead-time in stopped-flow measurements (**Figure S7B**), which confirms that interaction with both ligands does not lead to loop1 closure in these variants.

To obtain rate constants for association and dissociation kinetics of the reaction products AICAR and ImGP a dataset of 16 time traces was recorded by mixing excess of the ligand with limiting concentrations of HisF CouA or the binary complexes (HisFCouA*ImGP and His CouA*AICAR). A kinetic model describing ImGP/AICAR binding to HisF (**Figure S7C**) includes association and dissociation of the two ligands to the apo enzyme and to the respective binary complexes (*k*_1_ and *k*_-1_ for ImGP binding as well as *k*_2_ and *k*_-2_ for AICAR binding) and a conformational change (the closing of loop 1) to stabilize the ternary complex.

The stopped-flow datasets for wt-HisF (**Figure S8**) and variant HisF-F38A (**Figure S9**) were subjected to a global fitting analysis according to this kinetic model. The curves resulting from global fitting analysis are indicated by dashed lines. The determined values for the rate constants are summarized in **Table S4**. The rate constants obtained for the HisF-F38A variant in the global fitting analysis resemble those for wt-HisF. Importantly, the *K*_D_ values for the binding reactions that were calculated from the global fitting parameters roughly match the *K*_D_ values obtained in equilibrium titrations (*cf*. **Table S1** and **Table S4**). It should be emphasised that the binding model is a minimal model that accounts for key features of the experimental data. It could well be that the rate constants for binding of AICAR and ImGP to apo HisF and the HisF*ImGP/HisF*AICAR complex, respectively, differ. However, this cannot be better resolved with the stopped-flow datasets, as the binary enzyme-ligand complexes form within the dead time of the stopped-flow instrument. Importantly, the rates for the loop opening (*k*_-conf_) are higher for the ImGP:AICAR complex than for the PrFAR complex, which indicates that loop1 opens after the reaction to allow for product release.

### The motions of loop1 are not rate limiting in the kinetic mechanism of HisF

To discern which step in the catalytic mechanism is rate-determining for wt-HisF and HisF-G20P, HisF-F23A, and HisF-F38A, turnover kinetics under multiple turnover conditions were compared with turnover rates obtained with single turnover conditions. In the multiple turnover mode HisF was mixed with an excess of the substrate PrFAR and the turnover of PrFAR was monitored based on the decrease of absorption at 300 nm. Catalytic turnover of PrFAR by HisF occurs only in the presence of the second substrate ammonia. Therefore, we compared turnover traces in the presence of ammonia with control traces obtained in the absence of ammonia to discriminate absorption changes accompanying PrFAR turnover from signals stemming from binding or mixing reactions. For the reaction of wt-HisF a representative multiple turnover trace and the associated control trace are shown in Figure 8A. The corresponding data for the HisF variants are shown in **Figure S10A** (HisF-F38A), **Figure S11A** (HisF-F23A), and **Figure S12A** (HisF-G20P).

**Figure 8.**
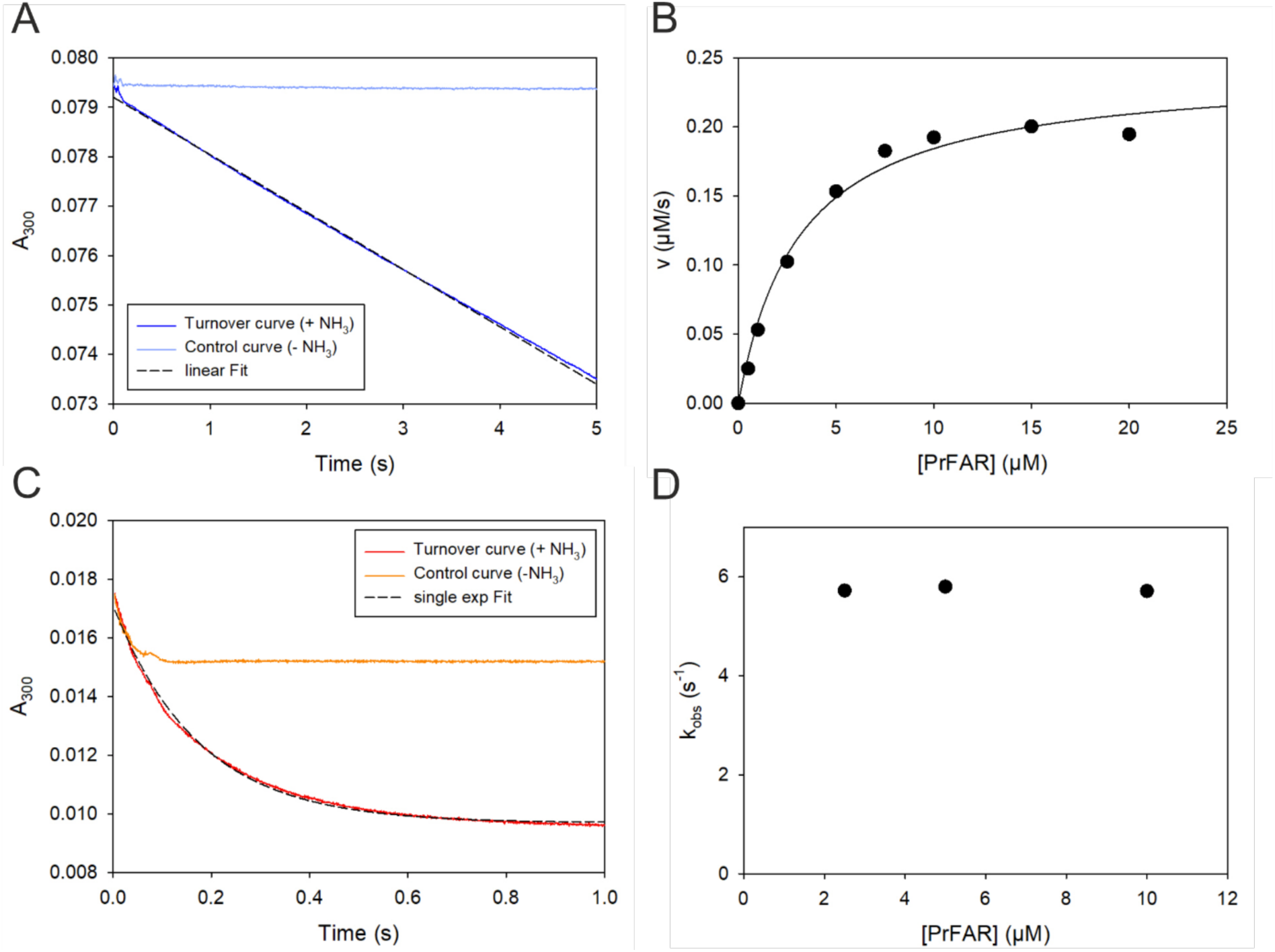
Multiple- and single-turnover kinetics of the wt-HisF reaction. **(A)** Representative transient monitoring PrFAR conversion in multiple turnover mode at 25°C after mixing 0.1 µM HisF with 10.0 µM PrFAR (final concentrations) in the presence of 100 mM ammonium acetate (turnover curve, blue line). A linear approximation of the steady-state phase (dashed line) yielded a turnover velocity of *v* = 0.192 µM s^-1^. The control curve (light blue line) shows the progress of the reaction in absence of ammonium acetate. **(B)** Plot of the turnover velocity *v vs.* the respective PrFAR concentration in multiple turnover experiments. *k*_cat_ and *K*M^PrFAR^-values were obtained by fitting to the Michaelis-Menten equation. **(C)** Representative transient monitoring PrFAR conversion in single turnover mode after mixing an excess of HisF (20 µM) with 10 µM PrFAR in the presence of 100 mM ammonium acetate (turnover curve, red line). The turnover curve was fit with a single exponential decay function (dashed line, 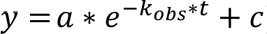). The control curve (orange line) shows the progress of the reaction in the absence of ammonium acetate. **(D)** Plot of the turnover rates, *k*_obs_, observed under single turnover conditions, *vs.* the respective PrFAR concentration. *k*_cat-_, *K*_M-_ and *k*_obs_-values are summarized in Table S5.

The time traces obtained under multiple turnover conditions showed a linear steady-state phase that is preceded by an exponential burst phase. The burst phase was observed also in the control curve in absence of ammonia. We attribute this burst phase to a mixing artifact of the stopped-flow instrument and this phase was not analyzed any further. Turnover velocities were deduced from a linear fit of the steady-state phase and were plotted as a function of the PrFAR concentration to obtain the *k*_cat_ and *K*M^PrFAR^ values for wt-HisF (Figure 8B) and HisF-F38A, HisF-F23A, and HisF-G20P (**Figures S10B – S12B**).

For measurements under single-turnover conditions the substrate PrFAR was saturated with enzyme so that all PrFAR molecules participate in the single turnover. The rate of turnover rate in that case is unaffected by product release and can be determined by fitting the change in the fluorescence over time to a single-exponential function. A representative single-turnover transient for wt-HisF is shown in Figure 8C. The corresponding data for the HisF variants are shown in **Figure S10C** (HisF-F38A), **Figure S11C** (HisF-F23A), and **Figure S12C** (HisF-G20P). Single-turnover rates *k*_obs_ determined from these exponential fits were independent of the applied PrFAR concentrations, both for wt-HisF (Figure 8D) and the variants HisF-F38A, HisF-F23A, and HisF-G20P (**Figure S10D-S12D**). The kinetic constants determined for the multiple and single turnover measurements are summarized in **Table S5**.

In summary, turnover rate measurements confirm that variant HisF-F38A is catalytically as active as wt-HisF, whereas the activities of the two loop1 variants HisF-F23A and HisF-G20P are significantly reduced. This deterioration of catalytic activity manifests mainly in *k*_cat_ values and single-turnover rates, which are reduced by three orders of magnitude, but is also expressed in a 2 to 5-fold increase of the *K*_M_ values. Remarkably, rate constants obtained in multiple and single turnover measurements have the same order of magnitude, implying that product release and associated conformational changes are not rate determining in the catalytic mechanism. Hence, it is concluded that the chemical step is rate-determining for catalysis by HisF. This is in contrast to other enzymes in this pathway, such as HisA and PriA, where loop motion is likely rate determining.^32^

## CONCLUSIONS

Due to their catalytic versatility and adaptability (βα)8-barrel enzymes have been successfully harnessed as scaffolds for enzyme design.^28, 66–67^ It is already becoming apparent that the inclusion of loop engineering into design strategies has an immense potential for the targeted engineering of substrate selectivity and catalytic activity.^68–70^ To do so, a comprehension of the conformational states of active-site loops and their significance for the catalytic mechanism is critical. Here, we have studied the importance of the flexible active-site loop1 for the kinetic mechanism of the (βα)8-barrel enzyme HisF. In several crystal structures, loop1 adopts a defined open conformation in the absence of substrates or a defined closed conformation when the binding partner HisH and substrates are bound in the active site.^40^ Furthermore, the NMR measurements presented here show that loop1, in the absence of substrates, adopts a highly flexible ensemble of detached conformations, which appear to be the predominant conformations in solution, and our molecular dynamics simulations indicate that the loop is conformationally plastic and capable of taking a range of conformational states, depending both on loop sequence and whether a ligand is bound to the active site or not (Figures 5).

To address the importance of the different loop conformations for catalytic turnover we shifted the conformation of loop1 through single point mutations. Subsequently we assessed the binding properties and activity of these variants through a combination of steady state and stopped-flow kinetics, X-ray crystallography, NMR spectroscopy and molecular dynamics simulations.

We established that loop1 in unliganded wt-HisF adopts both the open and detached conformations, where the substrate binding site is accessible, but without fully being able to access a catalytically competent closed conformation similar to that observed in the HisF/HisH complex (PDB ID: 7AC8^40^). After recruitment of the substrate, however, loop1 remodels and closes over the substrate binding pocket. In this closed conformation F23 in loop1 stacks onto the substrate. The formation of the catalytically important enzyme:substrate complex thus takes place in two steps: binding of substrate, followed by the closing of loop1 over the substrate.

In the HisF-F38A variant the open conformation of loop1 was slightly destabilized by removing an aromatic contact between this loop and the core of the enzyme. In the unliganded state, this led to a small shift from the open conformation towards the detached conformation. In the substrate-bound state, the HisF-F38A variant properly formed the closed conformation. In binding and activity assays, this variant was indistinguishable from wt-HisF. Based on that the equilibrium between the open and detached conformations does not play a rate limiting role in the catalytic cycle of the enzyme.

In the unliganded form of the HisF-G20P and HisF-F23A variants loop1 was slightly stabilized in the open conformation (G20P) or considerably shifted towards the detached conformation (F23A). As both variants interact with the substrate with a similar affinity this implies that the loop1 conformation in the apo state (open or detached) does not influence substrate recruitment. Nevertheless, both variants display a slightly reduced substrate binding affinity compared to the apo enzyme and, importantly, bind the substrate in a simple one step binding mechanism. In addition, NMR experiments reveal that these mutations impair the formation of the closed conformation. Consequently, the activity of these variants is reduced by three orders of magnitude compared to the wt-HisF protein.

Taken together, our data reveal a clear model that correlates conformational changes in loop1 with substrate turnover (Figure 9). In this model the formation of the closed loop1 conformation takes place after substrate recruitment and is essential for substrate turnover. After the cyclase reaction, which is the rate limiting step in the catalytic cycle, the closed loop conformation is destabilized, which facilitates product release.

**Figure 9.**
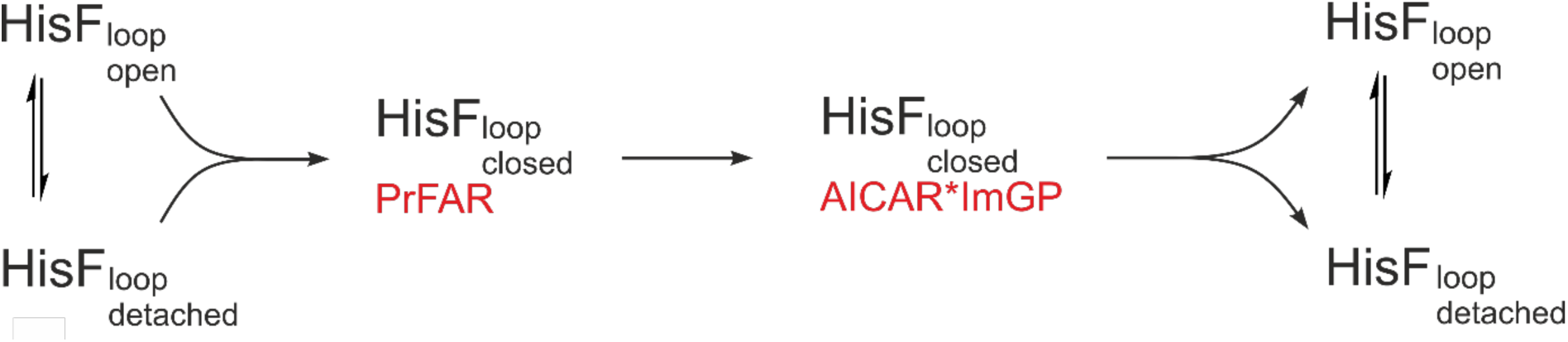
Model of the conformational changes of loop1 and their importance for the catalytic reaction.

It has been shown that ligand-gated loop motions in (βα)8-barrel enzymes differ in their magnitude (*e.g.,* number of loops involved) and dynamics, and affect different reaction steps of the catalytic mechanism. In the prototypical and well-studied TIM, which catalyzes the isomerization of glyceraldehyde-3-phosphate and dihydroxy-acetone phosphate in glycolysis, loop6 is a phosphate-gripper loop that moves ∼7 Å from a catalytically inactive open conformation to a catalytically active closed conformation upon substrate-binding.^71^ The conformational change occurs in concert with substantial internal rearrangement of the adjacent loop β7α7.^50, 72^ The most important consequence of the conformational change is the exclusion of solvent from the active site, reducing the dielectric constant in the surrounding of a catalytic glutamate residue and shifting its _p_*K*_a_ value in such a way that it can act as a general base.^73–74^ The movement of loop6 has long been interpreted as a rigid body movement, with the loop moving as a lid attached to two hinges.^33, 75–76^ However, more recent computational work indicates that loop6 is highly flexible, sampling multiple open distinct conformations that interconvert between each other, whereas the closed conformation falls in a very narrowly defined energy basin, and any deviation from this conformation has negative impact on catalytic turnover.^50^ Product release was identified as the rate-determining step in the biologically relevant reaction (conversion of dihydroxyacetone phosphate to *D*-glyceraldehyde 3-phosphate) and loop6 movement is necessary, among others, to release the product from the active-site.^76–77^

A similar role is played by loop movements in the catalytic mechanism of the (βα)8-barrel enzyme indole-3-glycerol phosphate synthase (IGPS, TrpC), which catalyzes the indole ring closure reaction during tryptophan biosynthesis.^78–80^ In IGPS, dynamics of the loop1, which houses a catalytically important Lys residue, are governed by competing interactions on the N- and C-terminal sides of the loop. Disrupting these interactions through amino acid substitutions quenches loop dynamics on the microsecond to millisecond timescales and slows down the dehydration step in the catalytic reaction.^81–82^ It seems that loop1 is maintained in a structurally dynamic state by the competing interactions, whereby the extent of loop mobility correlates with the rate-limiting step of the catalytic reaction and product release is rate-limiting at ambient temperature.^83^

In the case of HisA, PriA and TrpF, (βα)8-barrel enzymes that catalyze isomerization reaction in histidine and tryptophan biosynthesis, multiple active site loops undergo substantial ligand-gated conformational changes.^84–86^ Loop dynamics in HisA, PriA and TrpF is highly complex, with loop motion being at least partially rate limiting for substrate binding and being linked to substrate selectivity.^32^

L23oop1 motion in HisF differs from previous examples in so far as the chemical conversion itself, rather than substrate binding or product release, is rate-determining for the overall turnover reaction. A plausible reaction mechanism for the chemical conversion catalyzed by HisF has been proposed previously^36^ and involves the formation of two imine intermediates during acid-base catalysis. In the absence of structures of HisF bound to PrFAR or reaction intermediates, we can only speculate about how the induced fit facilitates the conversion reaction. It is reasonable to assume that the _p_*K*_a_ of the catalytic acid D11, which is very close to loop1, is increased by the generation of a hydrophobic environment stabilizing the protonated form of the aspartate side chain. When loop1 is in the closed conformation, F23 comes into close proximity of V18 and I52 and thereby create a hydrophobic cavity for the catalytic acid and shield it from solvent. A similar effect, the shielding of a catalytic aspartate residue through the closure of an active site loop has been described in TIM.^87^ In future work, this hypothesis may be substantiated by analyzing the protonation states of the catalytic residues D11 and D130 in HisF.

## MATERIALS AND METHODS

### Site-directed Mutagenesis

Point mutations were introduced into pET28a_HisF^88^ with a modified version of the protocol of the Phusion site-directed mutagenesis kit (Thermo Fisher Scientific) with HPSF-purified primers (BioSynth). To facilitate phosphorylation of the PCR product, T4 polynucleotide kinase was added during ligation. Mutagenesis was confirmed by Sanger sequencing (MicrosynthSeqlab). For the incorporation of the unnatural amino acid L-(7-hydroxycoumarin-4-yl)ethylglycine (CouA) at position 132, an amber stop codon mutation (TAG) was introduced into pET28a_HisF according to the protocol described above (pET28a_HisF_TAG).

### Protein Expression and Purification

All experiments were performed with *Thermotoga maritima* HisF (Uniprot ID: Q9X0C6) or HisF loop1 variants. Genes were expressed from modified pET vectors, encoding an N-terminal His6-tag followed by a TEV cleavage site, in *E. coli* BL21Gold (DE3) cells (Agilent Technologies). Expression was performed at 30°C overnight after induction with 1 mM Isopropyl β-D-1-thiogalactopyranoside (IPTG) at an OD600 of 0.6-0.8. Cells were harvested by centrifugation, resuspended in 50 mM Tris-HCl pH 7.5, 300 mM NaCl, 10 mM imidazole, and lysed by sonication. *E. coli* proteins were precipitated by a heat shock (15 min, 60°C) and removed by centrifugation. The supernatant was subjected to Ni-immobilized metal affinity chromatography (IMAC) (HisTrap FF Crude column, 5 ml, GE Healthcare). Proteins were eluted with a linear gradient of imidazole (10-500 mM). Fractions containing the protein of interest were identified by sodium dodecyl sulfate-polyacrylamide gel electrophoresis (SDS-PAGE) and pooled. Eluted proteins were digested with TEV protease at room temperature over night during dialysis against 50 mM Tris-HCl pH 7.5. TEV protease and non-cleaved protein was removed by IMAC (HisTrap FF Crude column, 5 ml, GE Healthcare) with a linear gradient of imidazole (0-500 mM). Fractions at low imidazole concentration containing the proteins of interest were identified by SDS-PAGE analysis, pooled, and further purified with a size-exclusion chromatography (SEC) column (Superdex 75 HiLoad26/260, GE Healthcare) by using 50 mM Tris-HCl pH 7.5 as the running buffer. Eluted protein fractions were checked by SDS-PAGE for > 90% purity, pooled, concentrated, and dripped into liquid nitrogen for storage at -80 °C.

For expression of HisF containing the unnatural amino acid CouA, pET28a_HisF_TAG was co-transformed with pEVOL_CouA, carrying the gene for the modified tyrosyl aminoacyl-tRNA synthethase from *M. janaschii*^57^ into *E. coli* BL21Gold (DE3). Cells were grown at 37°C in 6 L of LB medium until the OD600 reached 0.6-0.8. Cells were harvested by centrifugation and resuspended in 600 ml terrific broth (TB) medium. Bacterial growth at 37 °C was continued up to an OD600 of 10 and incorporation was induced by addition of 0.45 mM CouA and 0.02 % arabinose. Gene expression was induced by addition of 1 mM IPTG. Cultures were incubated overnight at 30 °C and the proteins were purified by nickel-affinity chromatography as described above.

The auxiliary enzymes HisA and HisE/IG from *T. maritima* were purified by standard methods from *E. coli* BL21-Gold cells (Agilent Technologies) that overexpressed the respective proteins.

### ProFAR/PrFAR Synthesis

The HisF ligands were synthesized enzymatically from 5-phospho-D-ribosyl α-1-pyrophosphate and adenosine triphosphate using the purified enzymes HisE/IG.^89^ The progress of the reaction was traced spectrophotometrically and the ProFAR product was purified using ion-exchange chromatography (POROS column; HQ 20, 10 ml, Applied Biosystems) using a linear gradient of 50 mM to 1 M ammonium acetate. ProFAR purity was examined through the absorbance ratio A290/A260 and the concentration was determined at a wavelength of 300 nm (ε_300_ = 6069 M^-1^ cm^-1^). PrFAR was synthesized from ProFAR with HisA from *T. maritima*. The product was purified using ion-exchange chromatography as described for ProFAR. Highly concentrated and >95% pure (A290/A260 = 1.1-1.2) fractions were unified, flash frozen in liquid nitrogen, and stored at -80°C.

### Limited Proteolysis

Proteolytic stability was tested at room temperature by incubating 10 µM HisF with 64 nM trypsin in 50 mM Tris/HCl pH 7.5. The reaction was stopped after different time intervals by adding one volume of 2x SDS-PAGE sample buffer and heating for 5 min at 95°C. The time course of proteolysis was followed on SDS-PAGE.

### Steady-state Enzyme Kinetics

The ammonia-dependent activity of HisF was measured by recording PrFAR turnover continuously at 300 nm [Δε_300_(PrFAR-AICAR) = 5637 M^-1^ cm^-1^] in 50 mM Tris/acetate pH 8.5 at 25 °C with a Jasco V650 UV-vis spectrophotometer. To determine *K*_M_^PrFAR^, 0.1-0.3 µM of wt-HisF or HisF loop1 variants were saturated with ammonia by adding 100 mM ammonium acetate (corresponding to 14.4 mM NH3 at pH 8.5). PrFAR (1-40 µM) was synthesized *in situ* from ProFAR, using a molar excess (0.5 µM) of HisA from *T. maritima* and converted by HisF to ImGP and AICAR. To ensure that ProFAR is completely turned over to PrFAR, the reaction mixture was incubated for at least 2 min before addition of wt-HisF or HisF loop1 variants. Enzyme activity was deduced from the initial slopes of the transition curves. Michaelis-Menten constants *K*_M_ and *k*_cat_ were determined by plotting the measured mean activity values and their standard error of the mean (SEM) of at least two technical replicates against the ProFAR concentration and fitting the data with the Michaelis-Menten equation (eqn. 1, 2):

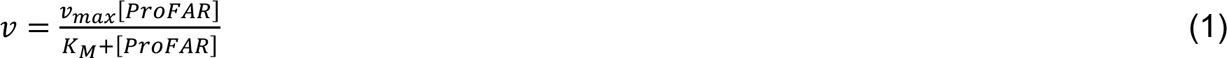

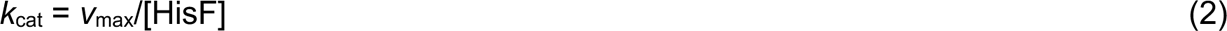

### Equilibrium Ligand Titrations

Fluorescence titrations of CouA-labeled HisF were performed at 25 °C in 50 mM Tris/acetate pH 8.5 in a Jasco FP-6500 spectrometer. CouA fluorescence emission was monitored at 451 nm with excitation at 367 nm. The substrate PrFAR was added stepwise from stock solutions in small volumes under constant stirring to a solution containing 0.2 µM CouA-labeled HisF and fluorescence emission was determined for each ligand concentration. Similarly, the product molecules AICAR or ImGP were titrated to 1.0 µM CouA-HisF (1.0 µM CouA-HisF/4.0 mM AICAR or 1.0 µM CouA-HisF/0.5 mM ImGP, respectively). Fluorescence values were corrected for dilution effects and the intrinsic fluorescence of ImGP. Fluorescence changes (ΔF) were plotted as a function of the ligand concentration and plots were fit to hyperbolic equations using SigmaPlot to obtain apparent *K*_D_ values (eqn. 3):

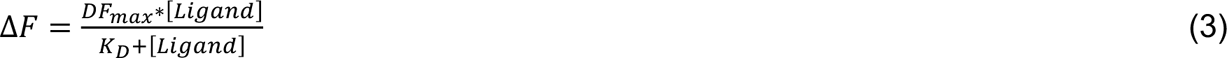

Given *K*_D_ values represent the average and standard error of at least two technical replicates.

### Stopped-flow Turnover and Ligand Binding Kinetics

Stopped-flow studies were performed at 25 °C using the SX20 stopped-flow instrument (Applied Photophysics). The instrument was equipped with a LED300 light source for absorbance measurements and a Me-Xe-Arc lamp for fluorescence measurements. At least five individual traces were recorded at each condition and averaged. Concentrations refer to final concentrations in the observation cell, unless otherwise specified.

Multiple- and single-turnover measurements were performed in 50 mM Tris/acetate (pH 8.5) and 100 mM ammonium acetate (turnover curves) or without ammonium acetate (control curves). For multiple-turnover measurements the A300 [△ε_300_(PrFAR-AICAR) = 5637 M^-1^ cm^-1^] was recorded over time after mixing a constant concentration of HisF (-wt/-F38A: 0.1 µM, - F23A: 0.5 µM or -G20P: 1.5 µM) with a molar excess of PrFAR (0.5 -50 µM) in a 1:1 volume ratio. Slopes were obtained by linear approximation of the steady-state part of the curves. The obtained turnover velocities (*v* = slope/(1 cm * 0.005637 µM^-1^ cm^-1^)) were replotted as a function of the PrFAR concentration and fitted with the Michaelis-Menten equation to obtain *k*_cat_ and *K*_M-_values. Single-turnover concentration series were measured by mixing excess HisF (20 µM) with different concentrations of PrFAR (2.5, 5.0 and 10.0 µM). Traces were fit with exponential decay functions 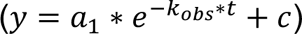. In the replot, *k*_obs_-values were plotted against associated PrFAR concentrations.

To study ligand binding kinetics, the change in fluorescence emission intensity of CouA-labeled HisF upon ligand binding was recorded over time in 50 mM Tris/acetate, pH 8.5 with an excitation wavelength of 367 nm and a 420 nm cut-off filter. For analysis of the PrFAR binding reaction, a constant concentration of CouA-labeled HisF (0.1 µM) was mixed with an excess of PrFAR (0.5-40 µM) in a 1:1 volume ratio to observe the binding reaction. Traces corresponding with wt-HisF and the HisF-F38A variant were fit to the sum of two exponential functions 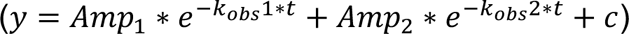, corresponding traces of variants HisF-F23A and HisF-G20P were fit to single exponential functions 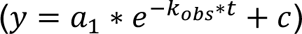.

To analyse the binding kinetics of the reaction products AICAR and ImGP, CouA-labeled HisF (0.05 µM) was mixed with an excess of AICAR (0.25 mM, 1.0 mM), an excess of ImGP (0.1 mM, 0.25 mM), or a mixture of the two molecules. In addition, the preformed binary complexes (0.05 µM HisF-CouA/1.25 mM AICAR, 0.05 µM HisF-CouA/0.4 mM ImGP) were mixed with the respective second product molecule (0.1-0.25 mM ImGP, 0.25-1.0 mM AICAR) to observe formation and dissociation of the ternary complex.

### Global Fitting Analysis

Sets of primary kinetic traces associated with the binding of ImGP and AICAR to HisF were fit globally to kinetic models using DynaFit (BioKin)^90^, which utilizes direct numerical integration to simulate experimental results. The script file for the global analysis of the binding reaction of wt-HisF is shown in the **Supplemental Methods**. Rate constants and the associated response coefficients were optimized iteratively in the global analysis. DynaFit features an error analysis functionality and model discrimination analysis, which was utilized to compare various kinetic models and to evaluate the quality of the fits.

### Protein Crystallization, X-ray Data Collection, and Structure Determination

For crystallization HisF was concentrated to 25 mg/ml and mixed 1:1 with the respective reservoir solution for hanging drop vapour diffusion crystallization. Crystals of wt-HisF were grown in previously determined conditions using Qiagen EasyXtal 15 well plates.^91^ HisF-F23A was crystallized in 1.2 M ammonium phosphate, using wt-HisF crystals for micro seeding. Crystals of HisF-G20P were obtained using a Morpheus II Screen (Molecular dimensions). Crystals were mounted on a nylon loop and flash frozen in liquid nitrogen without addition of cryoprotectants. Data sets were collected using synchrotron radiation from the Swiss Light Source (SLS), Switzerland at beamline PXIII and PXI. Data collection was done at cryogenic temperature (see **Table S6** for data collection and refinement statistics). Data were processed using XDS^92^, and the data quality was assessed using the program PHENIX^93^. Structures were determined by molecular replacement with MOLREP and programs within the CCP4isuite^94^ using PDB entry 1THF^48^ as the search model. Initial refinement was performed using REFMAC^95^. The model was further improved in several refinement rounds using automated restrained refinement with the program PHENIX^93^ and interactive modelling with Coot^96^.

### NMR Measurements

HisF used for ProFAR titration experiments was ^15^N-labeled. HisF used for backbone assignment and {^1^H}-^15^N hetNOE experiments were ^2^H, ^13^C, ^15^N-labeled. Isotope labelling was achieved by expression in *E. coli* BL21-CodonPlus (DE3) cells in M9 minimal medium. The M9 medium was H2O based and contained 0.5 g/L ^15^NH4Cl for expression of ^15^N-labeled protein and 2 g/l ^13^C -glucose for ^13^C-labeled samples. The M9 medium for the expression of ^2^H, ^13^C, ^15^N-labeled protein was D2O based and contained 0.5 g/L ^15^NH4Cl and 2 g/L ^2^H/^13^C-glucose. Protein expression was induced at an OD600 of 0.8 by addition of 1 mM IPTG to the medium and proteins were expressed overnight at 25 °C. Cells were harvested by centrifugation and lysed by sonication. *E. coli* proteins were precipitated by a heat shock (20 min, 70 °C). Precipitated proteins and cell debris were removed by centrifugation. Isotope-labeled HisF was purified by IMAC as described for non-labelled HisF. After TEV cleavage the ^2^H, ^13^C, ^15^N-labeled HisF was unfolded and refolded to exchange the ^2^H from the expression medium to ^1^H for the amide groups in the protein core. This was achieved by dialysis overnight at room temperature against 50 mM Arg/Glu pH 7.3, 25 mM HEPES, 5 M guanidinium chloride, 2 mM DTT for unfolding and subsequent dialysis overnight at room temperature against 50 mM Arg/Glu pH 7.3, 25 mM HEPES, 50 mM NaCl, 2 mM DTT. Afterwards the proteins were subjected to reverse IMAC. The flow through of this column was concentrated and purified by SEC (Superdex 75 HiLoad26/260, GE Healthcare) using NMR buffer (20 mM HEPES pH 7.3, 50 mM NaCl, 1 mM DTT) as running buffer.

NMR experiments were conducted at 30 °C in NMR buffer supplemented with 5 % (v/v) D2O on 600 and 800 MHz Bruker Avance Neo spectrometers equipped with N2 (600 MHz) or helium (800 MHz) cooled cryoprobes. NMR samples contained 100-200 µM ^15^N-labeled HisF for ProFAR titrations or 300-600 µM ^2^H, ^13^C, ^15^N-labeled HisF for {^1^H}-^15^N hetNOE and backbone assignment experiments. The previously published backbone assignments of wt-His^97^ that were obtained under different buffer conditions were transferred to our measurement conditions based on TROSY variants of 3D-HNCACB, 3D-HN(CO)CACB, 3D-HN(CA)CO and 3D-HNCO experiments.^98^ The same set of experiments was used to transfer the assignment from wt-HisF to the loop1 variants HisF-F23A, HisF-G20P, and HisF-F38A.

Due to the instability of ProFAR, which prevents the use of triple resonance spectra for backbone assignment, a selective unlabeling strategy was used for the assignment of F23 in the ProFAR-bound state (**Figure S13**): A ^15^N-labeled sample of HisF with non-labeled Tyr/Phe residues and a ^15^N^13^C-labeled sample with non-labeled Asn residues were prepared by addition of non-labeled amino acids (100 mg/L each) to ^15^N or ^15^N/^13^C H2O-M9 medium. Unlabeling of Tyr in addition to Phe was chosen due to isotope scrambling between the two amino acids. As F23 is the only Phe/Tyr succeeding an Asn it can be assigned by comparing ^1^H^15^N-TROSY spectra of the Tyr/Phe unlabeled sample and 2D ^1^H^15^N-HNCO spectra of the Asn unlabeled sample with fully labeled samples. Comparison of the ^1^H^15^N-TROSY spectra reveals all Tyr/Phe signals, whereas the comparison of the 2D ^1^H^15^N-HNCO spectra reveals all signals succeeding an Asn. Only the signal of F23 is missing in both spectra. As both, ^1^H^15^N-TROSY and ^1^H^15^N-HNCO spectra, can be recorded in a couple of hours this allows the unambiguous assignment of F23 even in the unstable ProFAR sample.

{^1^H}-^15^N hetNOE experiments were recorded using the pulse sequence from Lakomek et al.^99^ with a recovery delay of 1 s and a proton saturation time of 9 s. Spectra were processed with Topspin 4.0.2 or NMRPipe 9.6^100^. Spectra were analyzed with CARA^101^and integrated with NMRPipe^100^.

### Molecular Dynamics Simulations and Analysis

Molecular dynamics simulations with HisF were performed in the loop1 open and closed states, both with and without PrFAR bound to the active site. Due to the lack of a structure of HisF isolated from HisH in a loop closed conformation, all loop-closed simulations were performed by extracting wt-HisF coordinates from the crystal structure of the HisF/HisH complex (PDB ID: 7AC8^40^), with substrate PrFAR aligned with and replacing the crystallized substrate analogue ProFAR in the HisF active site. Loop open simulations were initiated from the loop-open structure of wt-HisF (PDB ID: 1THF^48^), with the introduction of a S21T reversion to match the other crystal structures used in this work. In both loop open and loop closed systems, starting structures of the HisF-F23A and HisF-F38A variants for simulation were constructed based on the corresponding wt-HisH crystal structure. In the case of the HisF-G20P variant, starting structures for loop closed simulations of this variant were generated based on the corresponding wt-HisF crystal structure, whereas the loop open simulations were initiated from the corresponding crystal structure of this HisF variant (PDB ID: 8S8R, this work). All manually generated mutant structures were created using PyMOL^102^ applying the “Mutagenesis” function. Rotamers were selected from the backbone-dependent rotamer library such as to eliminate structural clashes.

The resulting crystal structures were then prepared for simulations and equilibrated following a standard equilibration procedure, as described in detail in the **Supplemental Methods**. Once equilibrated, ten 1 µs production runs were performed for each system in an NPT ensemble (1 atm pressure and 300 K), resulting in 30 µs cumulative simulation time per variant (initiated from loop open and closed conformations for liganded systems but just open for unliganded ones), and 120 µs cumulative simulation time across all enzyme variants. Convergence of the simulations is shown in **Figures S14** – **S16**. Hydrogen atoms in all production simulations were scaled using hydrogen mass repartitioning^103^ allowing for a 4 fs simulation time step. Temperature and pressure were regulated using Langevin temperature control (collision frequency 1 ps^−1^), and a Berendsen barostat (1 ps pressure relaxation time). All simulations were performed using the AMBER ff14SB force field^104^, and the TIP3P water model^105^, using the CUDA-accelerated version of the Amber22 simulation package^106^. Further details of simulation setup, equilibration and analysis are provided as **Supplemental Methods**, and a data package containing simulation starting structures, snapshots from trajectories, representative input files and any non-standard simulation parameters has been uploaded to Zenodo for reproducibility and is available for download under a CC-BY license at DOI: 10.5281/zenodo.12211377.

## ASSOCIATED CONTENT

### Supporting Information

Supplemental Figures S1-S16, Supplemental Tables S1-S9, Supplemental Methods, Supplemental References.

## AUTHOR INFORMATION

### Corresponding Authors

**Shina Caroline Lynn Kamerlin** – *School of Chemistry and Biochemistry, Georgia Institute of Technology, Atlanta, GA-30332, USA*

orcid.org/0000-0002-3190-1173

**Remco Sprangers –** *Institute of Biophysics and Physical Biochemistry, Regensburg Center for Biochemistry, University of Regensburg, 93053 Regensburg, Germany*

orcid.org/0000-0001-7323-6047

**Reinhard Sterner –** *Institute of Biophysics and Physical Biochemistry, Regensburg Center for Biochemistry, University of Regensburg, 93053 Regensburg, Germany*

orcid.org/0000-0001-8177-8460;

### Authors

**Enrico Hupfeld** – *Technical University of Munich, Campus Straubing for Biotechnology and Sustainability. Chair of Chemistry of Biogenic Resources, 94315 Straubing, Germany*

**Sandra Schlee** – *Institute of Biophysics and Physical Biochemistry, Regensburg Center for Biochemistry, University of Regensburg, 93053 Regensburg, Germany* orcid.org/0009-0005-7728-9134

**Jan-Philip Wurm**, Bruker Biospin, Rudolf-Plank-Str.12, 76275 Ettlingen, Germany

**Chitra Rajendran** – *Institute of Biophysics and Physical Biochemistry, Regensburg Center for Biochemistry, University of Regensburg, 93053 Regensburg, Germany*

**Dariia Yehorova** – *School of Chemistry and Biochemistry, Georgia Institute of Technology, Atlanta, GA-30332, USA*

**Eva Vos** – *School of Chemistry and Biochemistry, Georgia Institute of Technology, Atlanta, GA-30332, USA*

**Dinesh Ravindra Raju** – *School of Chemistry and Biochemistry, Georgia Institute of Technology, Atlanta, GA-30332, USA*

### Author Contributions

E. H. conceptualized the project and performed enzymatic measurements and X-ray experiments. S. S. performed and analyzed transient kinetic measurements. J.P.W. performed, analyzed, and interpreted NMR experiments. C. R. performed and analyzed X-ray experiments. D.Y., E.V., and D. R. R. performed, analyzed, and interpreted MD simulations. E. H., S. S., and J. P. W. wrote the original draft. S.C.L.K., R.Sp. and R. St. supervised the project, acquired funding, and revised and edited the manuscript.

## Funding

This work was supported by a grant of the Deutsche Forschungsgemeinschaft (STE 891/11-2), and by the Swedish Research Council (grant number 2019-03499). The computational simulations and data handling were enabled by resources provided by the National Academic Infrastructure for Supercomputing in Sweden (NAISS) at Chalmers Centre for Computational Science and Engineering (C3SE), High Performance Computing Center North (HPC2N) and Uppsala Multidisciplinary Center for Advanced Computational Science (UPPMAX) partially funded by the Swedish Research Council through grant agreement no. 2022-06725 (SNIC 2022/3-2 and NAISS 2023/3-5). Additionally, this work used the Hive cluster, which is supported by the National Science Foundation under grant number 1828187 and was supported in part through research cyberinfrastructure resources and services provided by the Partnership for an Advanced Computing Environment (PACE) at the Georgia Institute of Technology, Atlanta, Georgia, USA. Further simulations were performed on the Theta cluster at the Argonne Leadership Computing Facility (ALCF), through a Director’s Discretionary award.

### Notes

The authors declare no competing financial interest.

## ACKNOWLEDGMENTS

The authors thank Jeannette Ueckert and Sabine Laberer for excellent technical assistance, Johanna Stoefl for support in the production of proteins for NMR studies, as well as Frank Raushel, Matthias Wilmanns, and Sihyun Sung for critical reading of the manuscript and for fruitful discussions.

## ABBREVIATIONS

AICAR, 5-aminoimidazol-4-carboxamidribotide; HisF, cyclase subunit of ImGPS (used for HisF from T. maritima); HisH, glutaminase subunit of ImGPS; IGPS, indole glycerol phosphate synthase; ImGP, imidazole glycerol phosphate; ImGPS, imidazole glycerol phosphate synthase, PDB, Protein Data Bank; PrFAR, *N*’-[(5’-phosphoribulosyl)formimino]-5-amino-imidazole-4-carboxamide ribonucleotide (HisF substrate); ProFAR, *N*’-[(5’-phosphoribosyl)formimino]-5-amino-imidazole-4-carboxamide ribonucleotide (HisF substrate analogue); TIM, triose phosphate isomerase; wt, wild type.

