## Supplementary Information for "Conformational modulation of a mobile loop controls catalysis in the (βα)_8_-barrel enzyme of histidine biosynthesis HisF"

[#]These two authors contributed equally

### Table of Contents

S1. Supplemental Figures ........................................................................................... S3

S2. Supplemental Tables ........................................................................................... S18

S3. Supplemental Methods ......…………………………………………………….…………… S24

S4. Supplemental References ……………………………………………………………….. S27

#### S1. SUPPLEMENTAL FIGURES

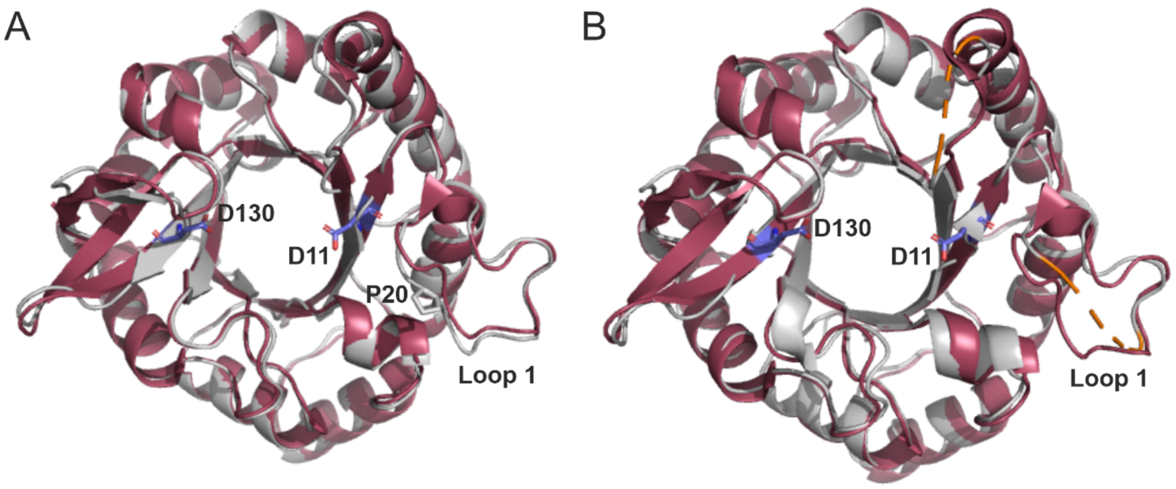


**Figure S1: Crystal structures of HisF-G20P and HisF-F23A in comparison with wt-HisF.**
Alignment of the crystal structure of wt-HisF (PDB-code 1VH7[1], red) with the structures of (**A**) HisF-G20P (PDB-code 8S8R, grey) and (**B**) HisF-F23A (PDB-code 8S8S, grey). The catalytic residues D11 and D130 are shown as blue sticks. Introduction of P20 leads to a slight distortion of the loop conformation. In the structure of variant HisF-F23A electron density for residues 20-24 (loop1) and residues 53-54 (loop2) is missing (indicated by dashed orange lines).

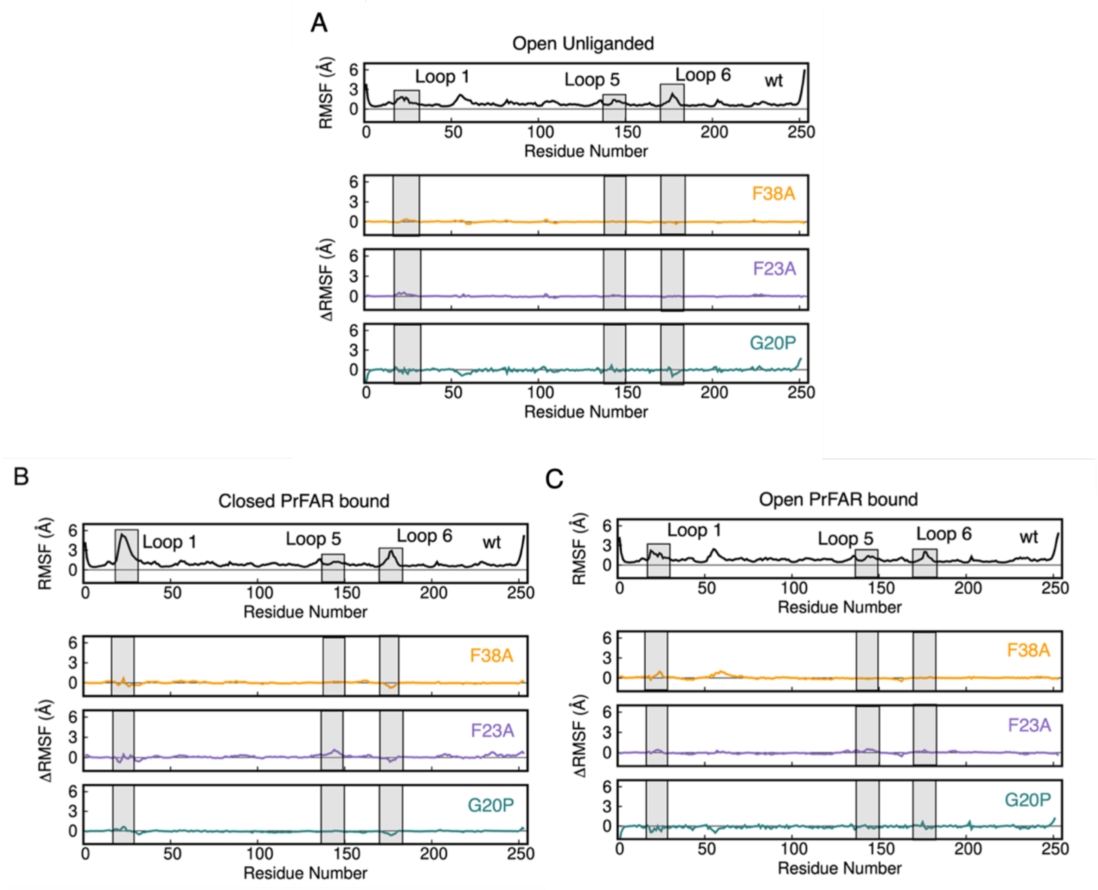


**Figure S2: Root mean square fluctuations (RMSF, Å) of the protein C$_\alpha$-atoms during molecular dynamics simulations of unliganded and PrFAR-bound HisF.**
Shown in each panel are the absolute C$_\alpha$-atom RMSF during simulations of wt-HisF (black), as well as the relative RMSF (ΔRMSF) between wt-HisF and each of the HisF-F38A (orange), HisF-F23A (purple), and HisF-G20P (green) variants. Shown here is data from simulations initiated from the (**A**) open unliganded, (**B**) closed PrFAR-bound, and (**C**) open PrFAR-bound states of each enzyme. Data were collected every 10 ps from 5 individual replicas of 1 μs length each.

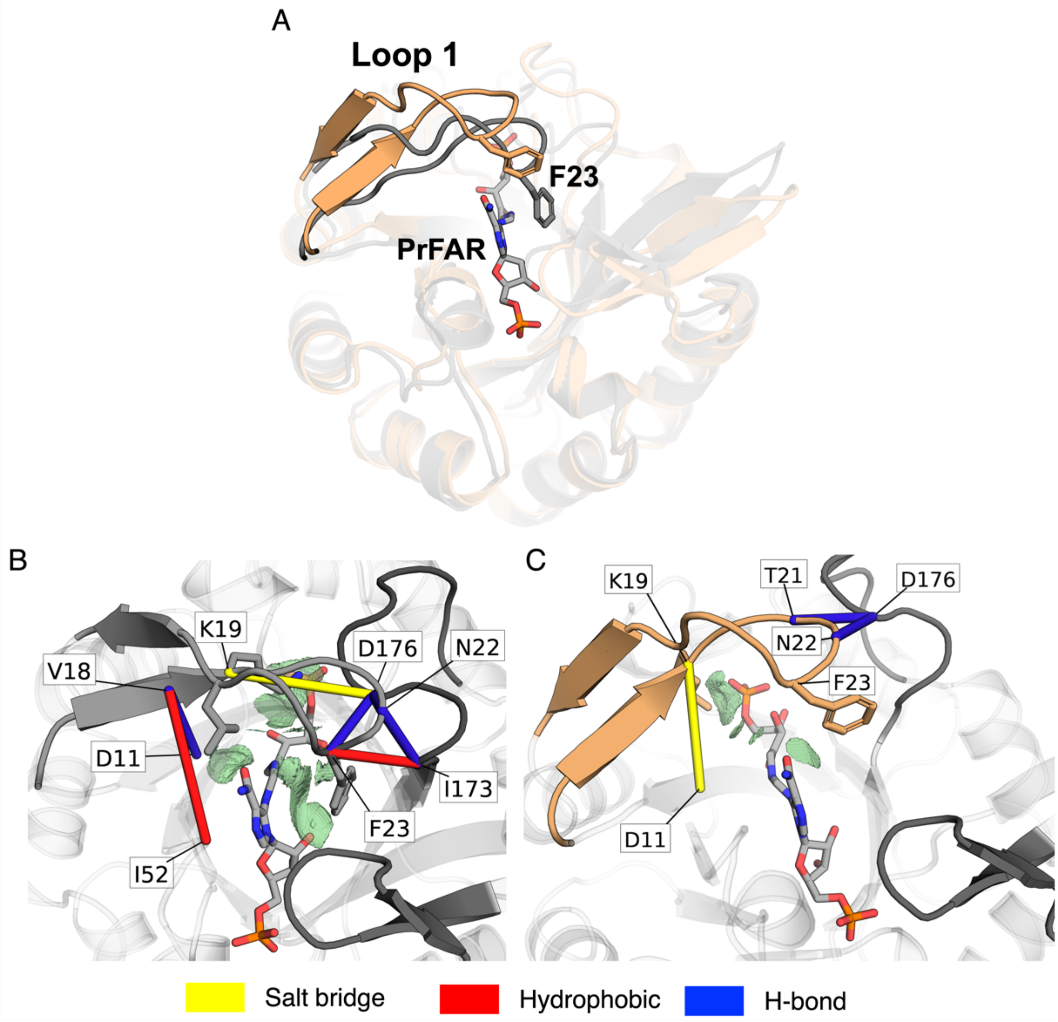


**Figure S3: Representative structures and interactions for the different closed conformations of wt-HisF from the X-ray structure and from simulations.**
(**A**) Structural comparison between the loop1-closed conformation observed in the HisF/HisH complex (grey, PDB ID:7AC8[2], chain E) in its ProFAR-bound state and the closed conformation obtained from the simulations of HisF in its PrFAR-bound state (tan). As can be seen from these structures, while we obtain loop1 conformations that have structural similarity to the closed conformation observed in the HisF/HisH complex, the loop is displaced compared to the conformation in the crystal structure. The bottom panels (**B**-**C**) show the results of interaction analysis performed using Key Interaction Networks[3] (KIN) on the (**B**) starting X-ray structure of HisF taken from the HisF/HisH complex, and (**C**) on closed conformations of loop1 from simulations of HisF in its PrFAR-bound states. The stick colors correspond to the different types of non-covalent interactions identified by KIN, where yellow represents salt bridges, red represents hydrophobic interactions, and blue represents H-bonding interactions. Interactions between loop1 and PrFAR were calculated using NCIplot[4] and are represented by the green surface of the electronic density.

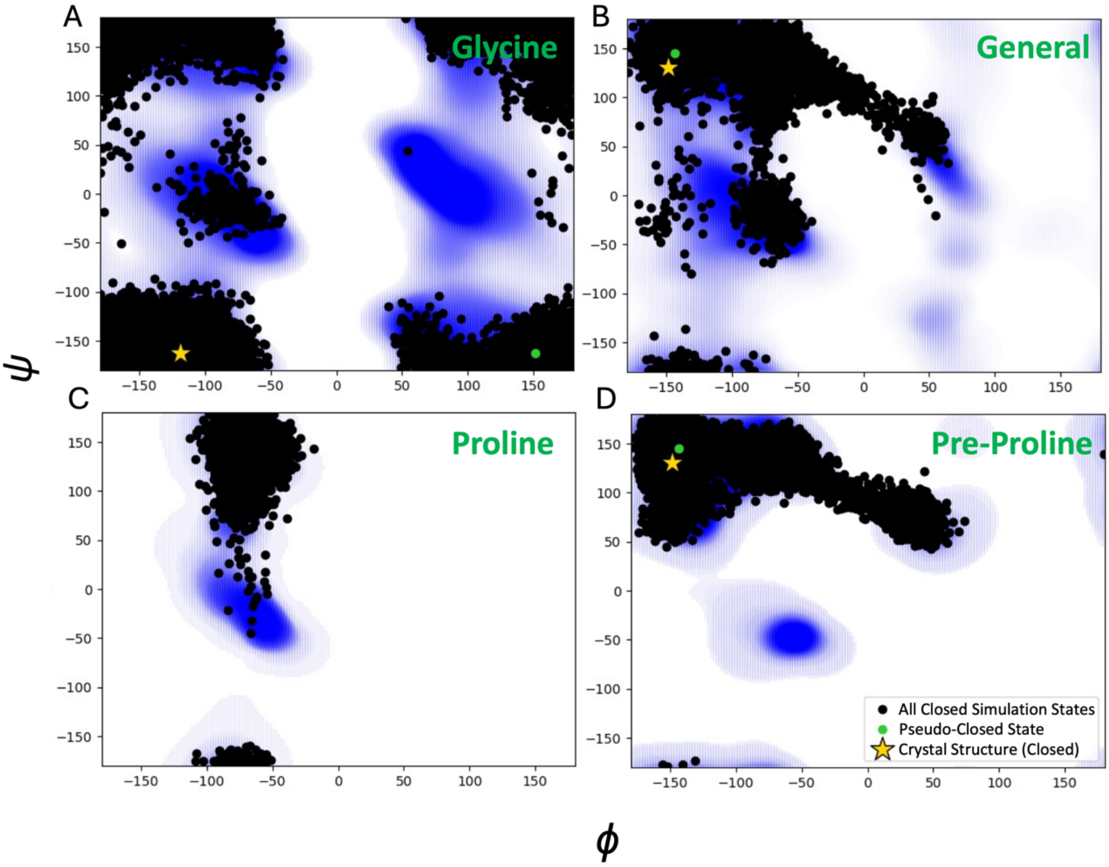


**Figure S4: Comparison of Ramachandran plots of positions 19 and 20 in wt-HisF and HisF-G20P.** Shown here are dihedral angles observed in residues K19 and G20/P20 of wt-HisF and the HisF-G20P variant overlayed over the Ramachandran plots that corresponds to the identity of the residue. Angles from the closed state simulations are plotted in black, while possible angles according to Ramachandran plot are shown in blue. Dihedral angles of the residues G20 and K19 for the closed crystal structure and a representative closed conformation are indicated by stars and green circles, respectively. This figure compares angles observed in (**A**) G20 of wt-HisF overlayed on glycine Ramachandran plot, (**B**) K19 in wt-HisF overlayed on general case Ramachandran plot, (**C**) P20 of HisF-G20P on the proline Ramachandran plot and (**D**) K19 in HisF-G20P on the pre-proline Ramachandran plot.

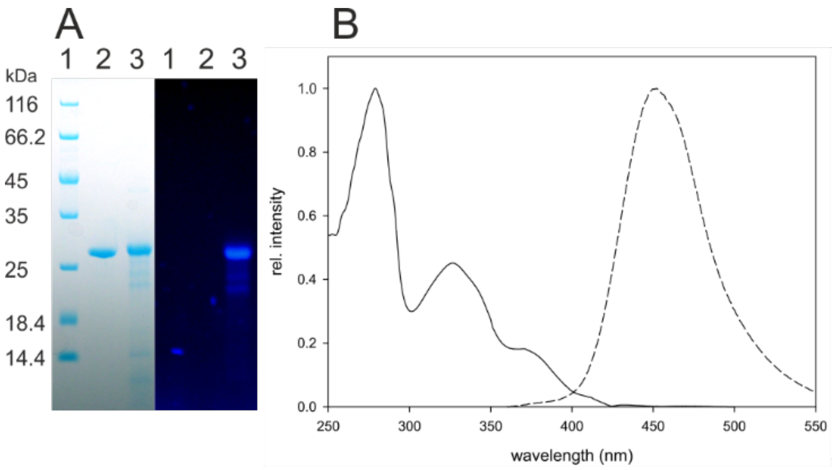


**Figure S5: Spectral properties of wt-HisF-K132CouA.**
(**A**) SDS-PAGE gel after Coomassie staining and illumination with white light (left) and UV light (333 nm) (right). Lane 1: Molecular weight standard, lane 2: non-labelled wt-HisF, lane 3: wt-HisF-K132CouA. Under UV light the typical blue fluorescence of CouA becomes visible. (**B**) The absorbance spectrum of HisF-K132CouA (solid line) encompasses peaks at 330 and 370 nm, characteristic for CouA, in addition to the protein absorption maximum at 280 nm. The fluorescence emission spectrum (dashed line) shows a maximum at 450 nm, inherent to CouA, after excitation at 370 nm. Intensities of the spectra have been normalized to the maximal signal.

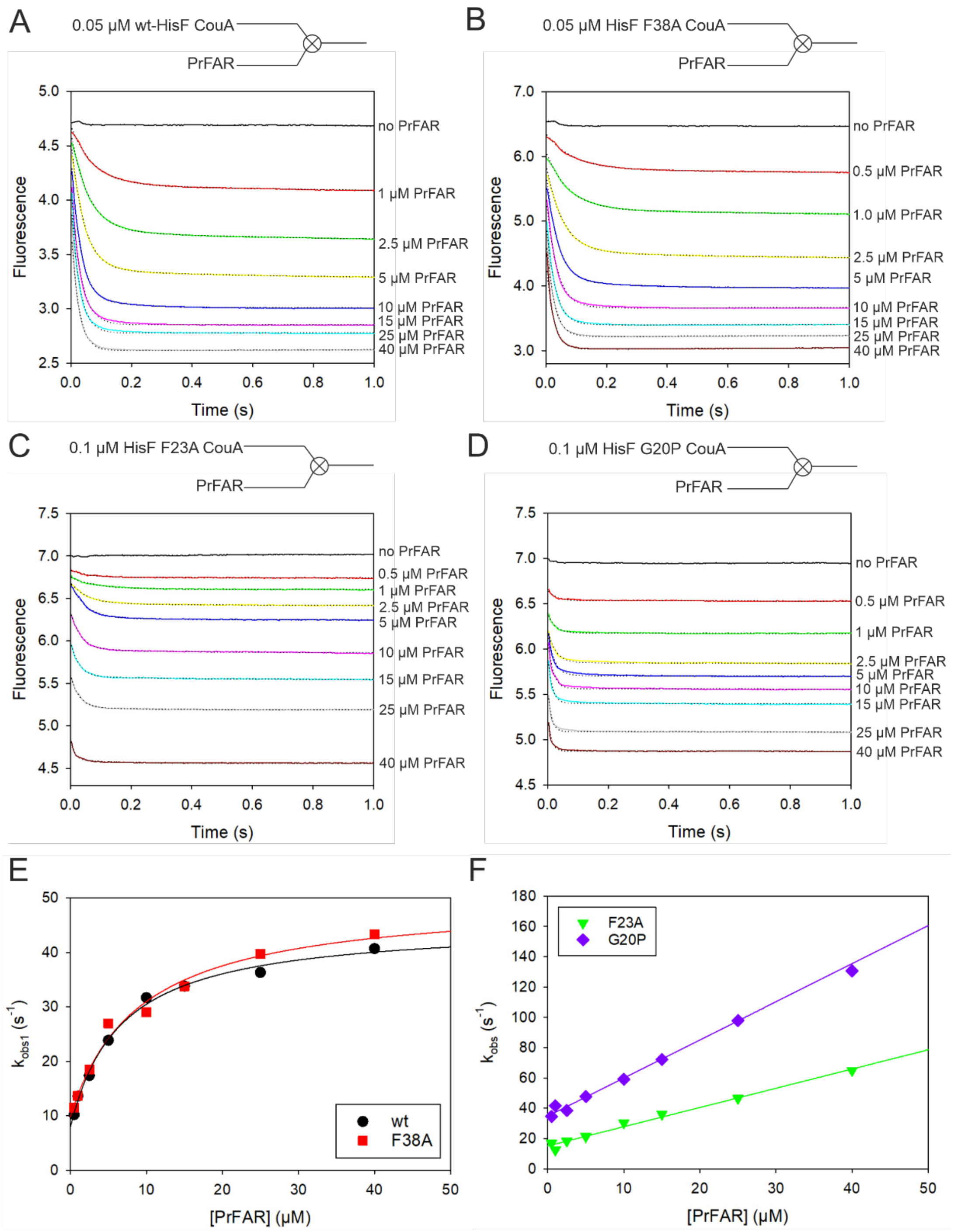


**Figure S6: Kinetics of PrFAR binding to HisF-CouA monitored by stopped-flow measurements.** (**A**) Time traces (coloured lines) recorded after mixing 0.05 µM wt-HisF with excess PrFAR at 25°C (Ex. 367 nm, Em cut-off: 400 nm). Time traces were fit to double exponential equations (dotted black lines). (**B**) Time traces recorded after mixing 0.1 µM HisF-F38A with excess PrFAR. Time traces were fit to double exponential equations (dotted black lines) (**C**) Time traces recorded after mixing 0.05 µM HisF-F23A with excess PrFAR. Time traces were fit to single exponential equations (dotted black lines). (**D**) Time traces recorded after mixing 0.1 µM HisF-G20P with excess PrFAR. Time traces were fit to single exponential equations (dotted black lines). (**E**) $k_{obs}$1-values associated with binding of PrFAR to wt-HisF-CouA and variant HisF-F38A-CouA were plotted *vs*. PrFAR concentration. The resulting plots show hyperbolic curves and were fitted to the equation $k_{obs}1 = k_{-conf} + k_{conf}$ *[PrFAR]/$K_{D1}$ + [PrFAR]) according to an induced fit model. (F) $k_{obs}$1-values associated with binding of PrFAR to variants HisF-F23A-CouA and HisF-G20P-CouA were plotted *vs*. PrFAR concentration. The resulting plots show linear dependencies and were fitted to linear equations ($k_{obs} = k_{-1} + k_1$ * [PrFAR]) according to a simple binding model. Numerical values obtained by curve fitting are listed in **Table S3**.

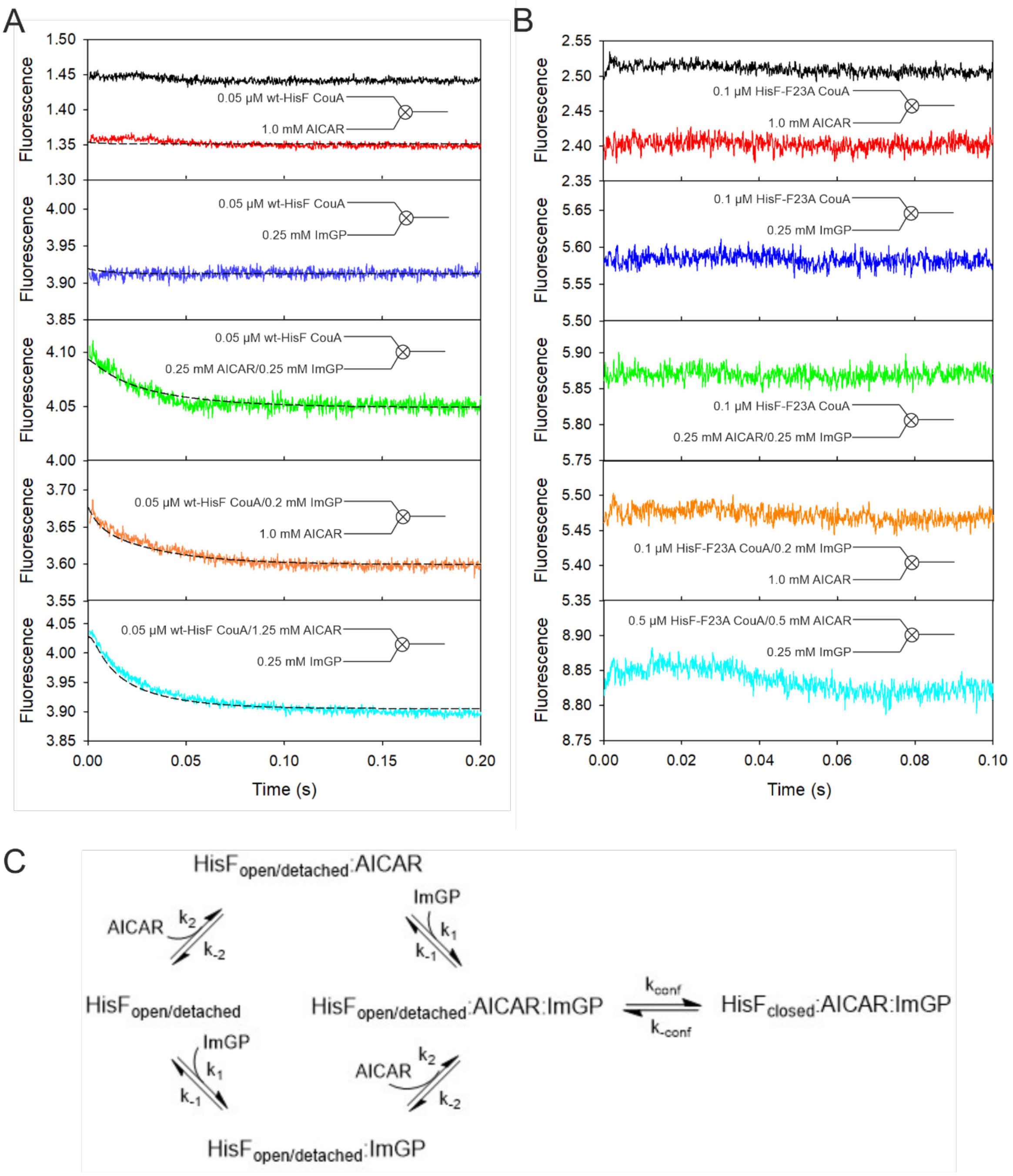


**Figure S7: Kinetics of ImGP and AICAR binding to HisF-CouA monitored by stopped-flow measurements.**

(**A**) Time traces were recorded at 25°C after mixing equal volumes of wt-HisF and buffer (black), AICAR (red), ImGP (blue) or AICAR/ImGP (green). In the same way, the pre-formed binary complexes wt-HisF CouA/ImGP (orange) and wt-HisF CouA/AICAR (cyan) were mixed with the respective second reaction product AICAR or ImGP and the corresponding time curves were recorded. The formation of the binary complexes is completed within the dead-time of the stopped-flow instrument, while formation of the ternary complex $HisF_{closed}$:AICAR:ImGP is associated with a decrease in fluorescence with an observed rate constant $k_{obs}$ in the range of 50 $s^{-1}$. (**B**) Time traces obtained after mixing HisF-F23A with AICAR and ImGP. The formation of both, the binary and ternary complexes is completed within the instrument dead time. Specified concentrations refer to final concentrations in the observation cell. (**C**) Kinetic model for the interaction of the reaction products ImGP and AICAR with wt-HisF and HisF-F38A including a conformational change upon formation of the ternary complex.

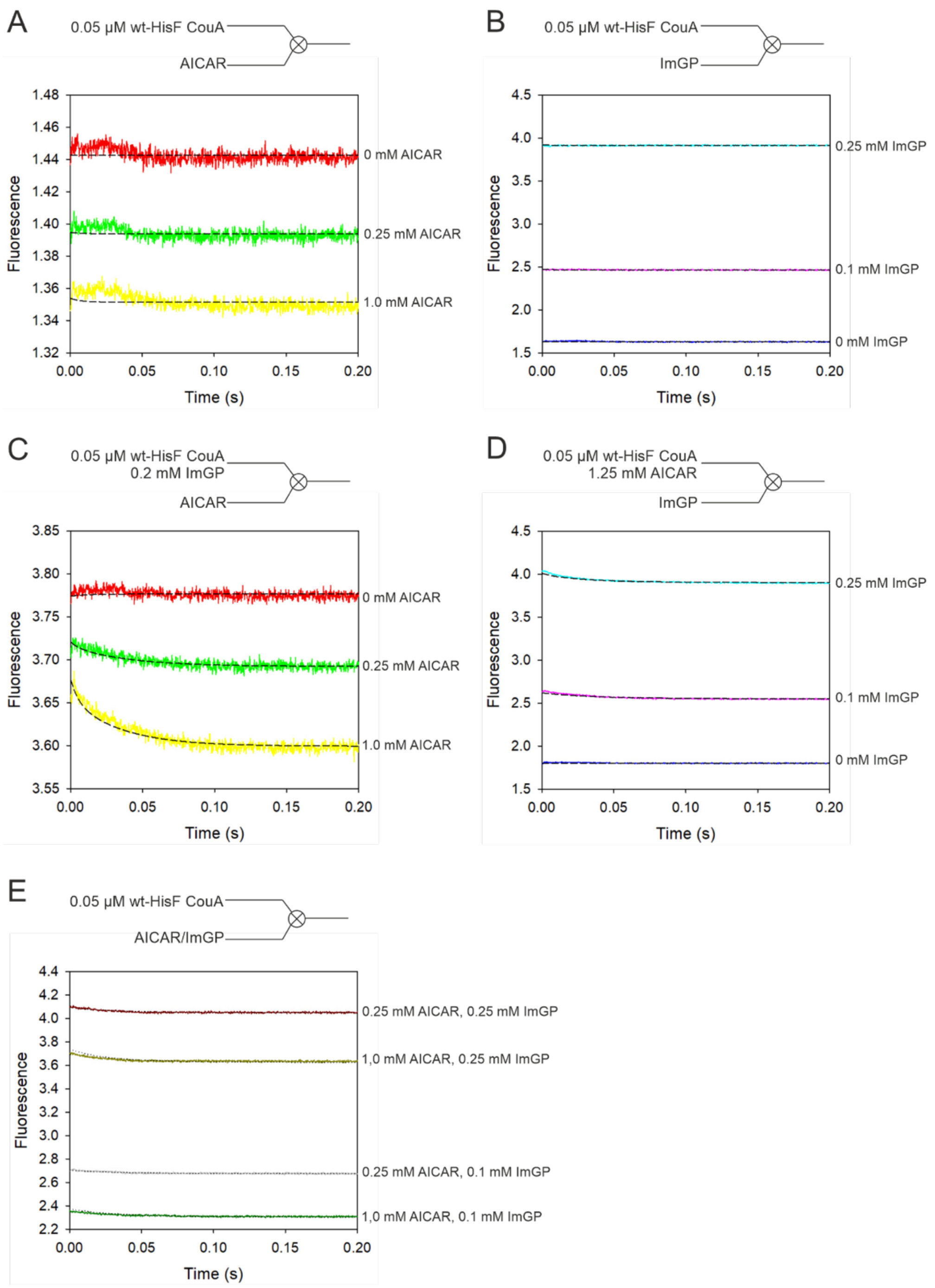


**Figure S8: Analysis of the binding reaction of AICAR and ImGP to wt-HisF-CouA.**
Kinetic traces monitoring the fluorescence changes at 25°C associated with binding of ImGP/AICAR to HisF are shown (Ex. 367 nm, Em cut-off: 420 nm). A dataset of 16 time traces was recorded by mixing limiting concentrations of HisF or the binary complexes (HisF-*ImGP and HisF*AICAR) with an excess of ligand. A kinetic model corresponding to the scheme in **Figure S8C** was fit to the stopped-flow transients, dashed lines represent the best fit after optimization of parameters in global fitting procedures. The fit parameters are summarized in **Table S4**. Concentrations denote final concentrations in the observation cell.
(**A**) Time traces obtained after mixing HisF with AICAR.
(**B**) Time traces obtained after mixing HisF with ImGP.
(**C**) Time traces obtained after mixing the preformed HisF*ImGP complex with AICAR.
(**D**) Time traces obtained after mixing the preformed HisF*AICAR complex with ImGP.
(**E**) Time traces obtained after mixing HisF with combinations of AICAR/ImGP.

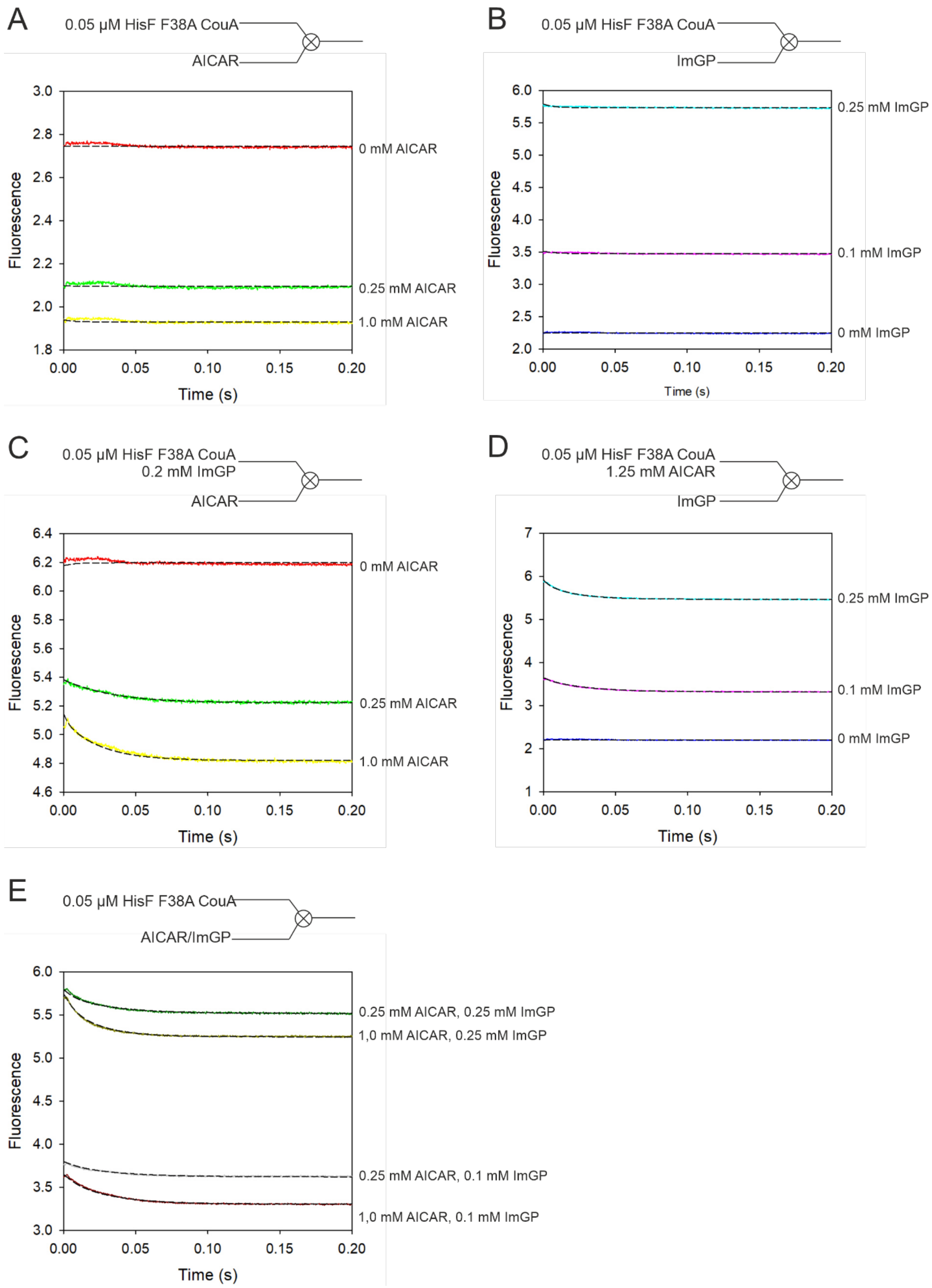


**Figure S9: Analysis of the binding reaction of AICAR and ImGP to HisF-F38A-CouA.**
Kinetic traces monitoring the fluorescence changes at 25°C associated with binding of ImGP/AICAR to HisF-F38A are shown (Ex. 367 nm, Em cut-off: 420 nm). A dataset of 16 time traces was recorded by mixing limiting concentrations of HisF-F38A or the binary complexes (HisF-F38A-*ImGP and HisF-F38A-*AICAR) with an excess of ligand. A kinetic model corresponding to the scheme in **Figure S8C** was fit to the stopped-flow transients, dashed lines represent the best fit after optimization of parameters in global fitting procedures. The fit parameters are summarized in **Table S4**. Concentrations denote final concentrations in the observation cell.
(**A**) Time traces obtained after mixing HisF-F38A with AICAR.
(**B**) Time traces obtained after mixing HisF-F38A with ImGP.
(**C**) Time traces obtained after mixing the preformed HisF-F38A*ImGP complex with AICAR.
(**D**) Time traces obtained after mixing the preformed HisF-F38A-*AICAR complex with ImGP.
(**E**) Time traces obtained after mixing HisF-F38A with combinations of AICAR/ImGP.

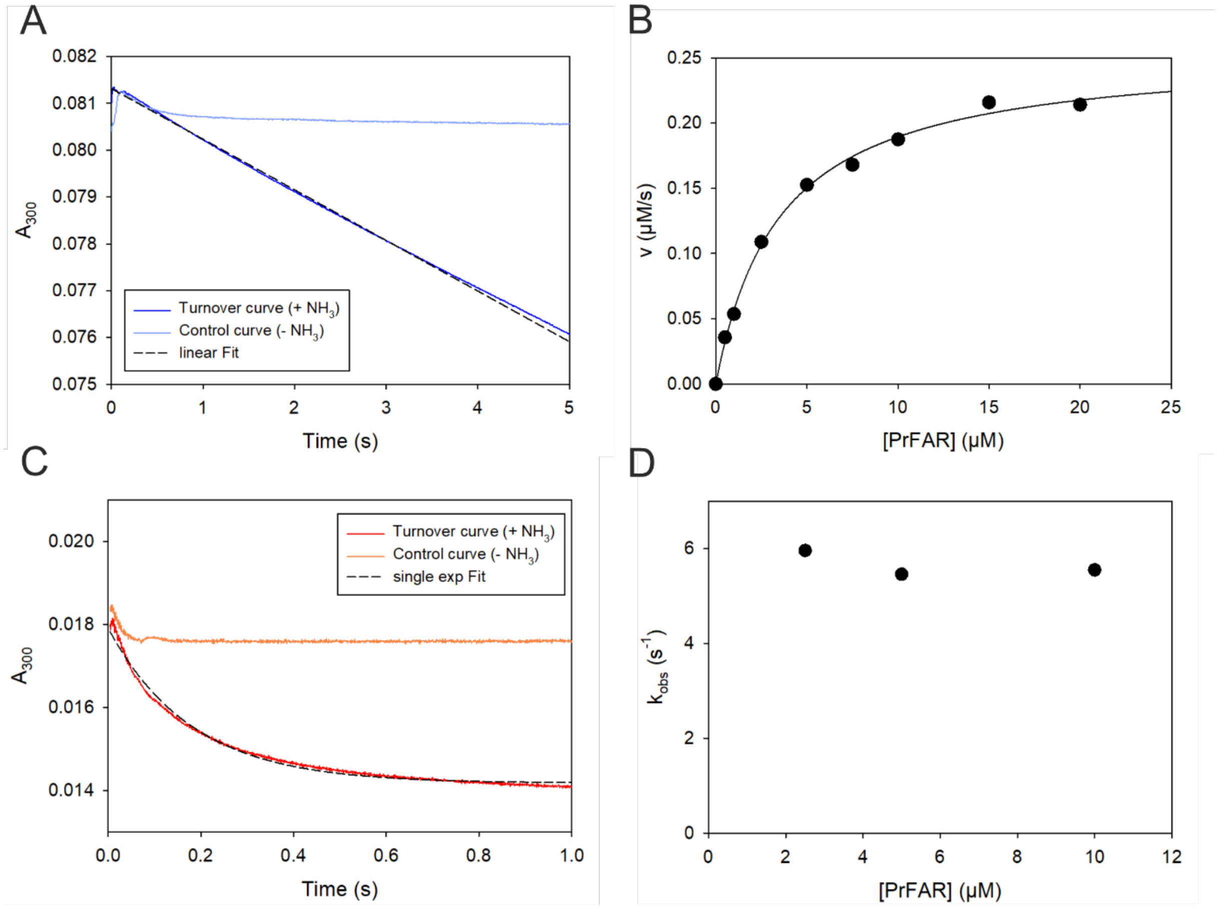


**Figure S10. Multiple- and single turnover kinetics of HisF-F38A.**
(**A**) A representative transient is shown monitoring PrFAR turnover in multiple turnover mode at 25°C after mixing 0.1 µM HisF-F38A with 10.0 µM PrFAR in presence of ammonium acetate (turnover curve, blue line). A linear approximation of the steady-state phase (dashed line) yielded a turnover velocity of $v$ = 0.188 µM $s^{-1}$. The control curve (light blue line) shows the progress of the reaction in absence of ammonium acetate. (**B**) Plot of the turnover velocity $v$ vs. the respective PrFAR concentration in multiple turnover experiments. The data were fit to the Michaelis-Menten equation. (**C**) A representative transient is shown monitoring PrFAR turnover in single turnover mode after mixing an excess of HisF-F38A (20 µM) with 10 µM PrFAR in the presence of 100 mM ammonium acetate (turnover curve, red line). The turnover curve was fit with a single exponential decay function (dashed line, $y = a * e^{-k_{obs}*t} + c$). The control curve (orange line) shows the progress of the reaction in the absence of ammonium acetate. (**D**) Plot of the turnover rates, $k_{obs}$, observed under single turnover conditions, *vs.* the respective PrFAR concentration. $k_{cat}$-, $K_M$- and $k_{obs}$-values are summarized in **Table S5.**

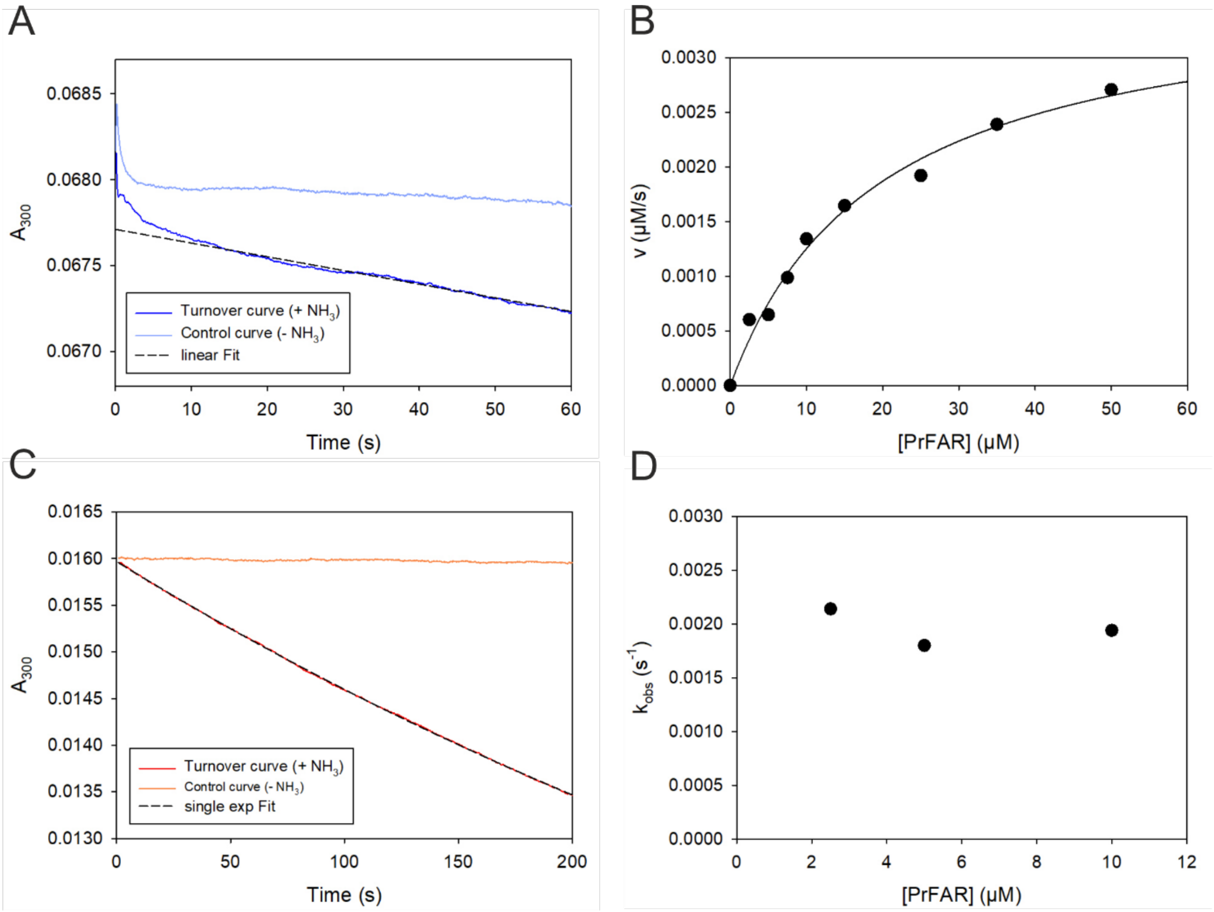


**Figure S11: Multiple- and single turnover kinetics of HisF-F23A.**
(**A**) A representative transient is shown monitoring PrFAR turnover in multiple turnover mode at 25°C after mixing 0.5 µM HisF-F23A with 10.0 µM PrFAR in presence of ammonium acetate (turnover curve, blue line). A linear approximation of the steady-state phase (20-60 s, dashed line) yielded a turnover velocity of $v$ = 1.34 x $10^{-3}$ µM $s^{-1}$. The control curve (light blue line) shows the progress of the reaction in absence of ammonium acetate. (**B**) Plot of the turnover velocity $v$ *vs.* the respective PrFAR concentration in multiple turnover experiments. The data were fit to the Michaelis-Menten equation. (**C**) A representative transient is shown monitoring PrFAR turnover in single turnover mode after mixing an excess of HisF-F23A (20 µM) with 10 µM PrFAR in the presence of ammonium acetate (turnover curve, red line). The turnover curve was fit with a single exponential decay function (dashed line, $y = a * e^{-k_{obs} * t} + c$). The control curve (orange line) shows the progress of the reaction in the absence of ammonium acetate. (**D**) Plot of the turnover rates, $k_{obs}$, observed under single turnover conditions, *vs.* the respective PrFAR concentration. $k_{cat}$-, $K_M$- and $k_{obs}$-values are summarized in **Table S5**.

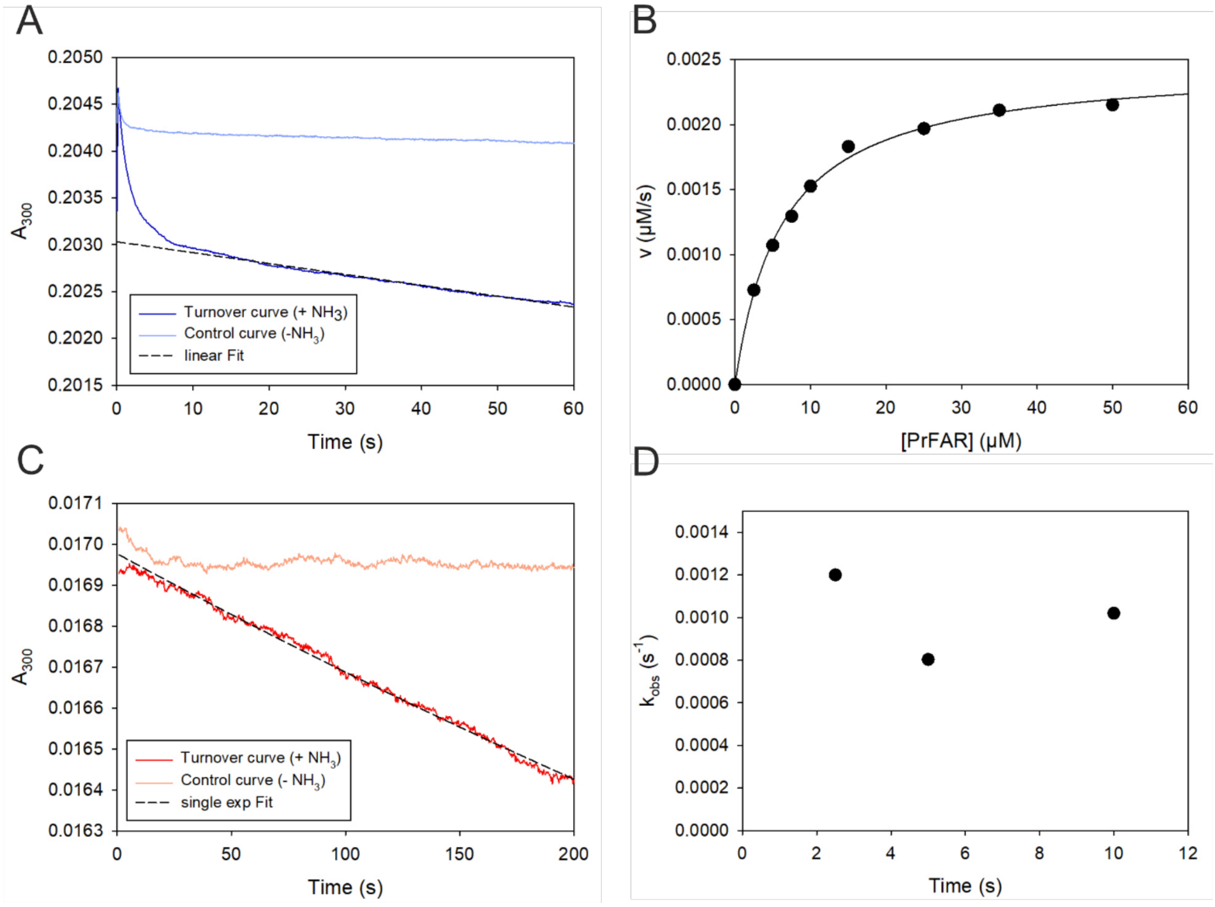


**Figure S12: Multiple- and single turnover kinetics of HisF-G20P.**
(**A**) A representative transient is shown monitoring PrFAR turnover in multiple turnover mode at 25°C after mixing 1.5 µM HisF-G20P with 25.0 µM PrFAR in the presence of 100 mM ammonium acetate (turnover curve, blue line). A linear approximation of the steady-state phase (dashed line) yielded a turnover velocity of $v$ = 2.1 x $10^{-3}$ µM $s^{-1}$. The control curve (light blue line) shows the progress of the reaction in absence of ammonium acetate. (**B**) Plot of the turnover velocity $v$ *vs.* the respective PrFAR concentration in multiple turnover experiments. The data were fit to the Michaelis-Menten equation. (**C**) A representative transient is shown monitoring PrFAR turnover in single turnover mode at 25°C after mixing an excess of HisF-G20P (20 µM) with 10 µM PrFAR in the presence of 100 mM ammonium acetate (turnover curve, red line). The turnover curve was fit with a single exponential decay function (dashed line, $y = a * e^{-k_{obs}*t} + c$). The control curve (orange line) shows the progress of the reaction in absence of ammonium acetate. (**D**) Plot of the turnover rates, $k_{obs}$, observed under single turnover conditions, *vs.* the respective PrFAR concentration. $k_{cat}$-, $K_M$- and $k_{obs}$-values are summarized in **Table S5**.

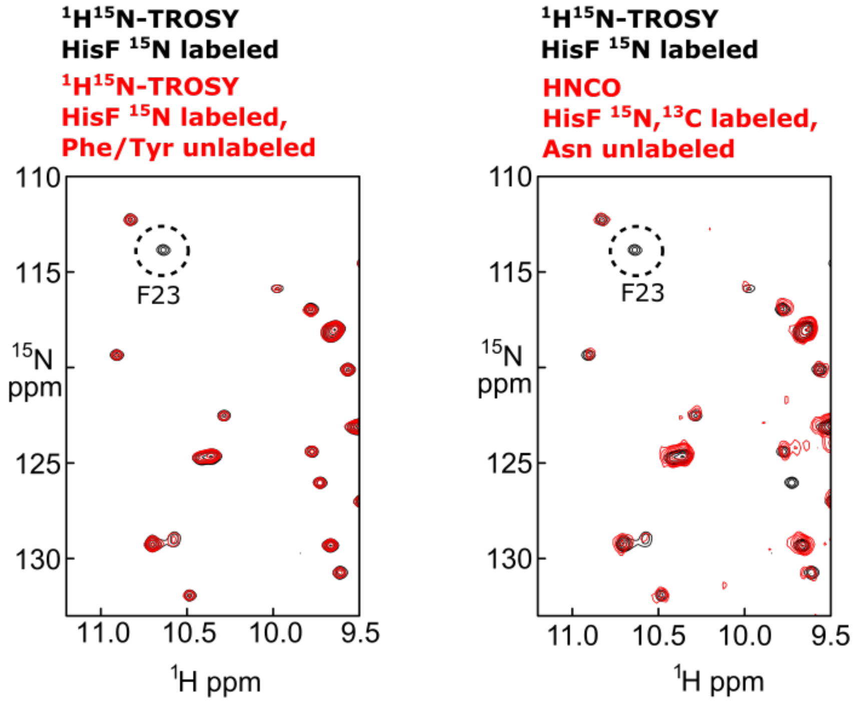


**Figure S13: Assignment of the F23 signal in the ProFAR-bound state of wt-HisF.**
(Left) Overlay of $^{1}H^{15}N$-TROSY spectra of fully $^{15}N$-labeled HisF in the presence of ProFAR (black) and of $^{15}N$-labeled HisF, where the Phe and Tyr residues are unlabelled, in the presence of ProFAR (red). The signal of F23 is indicated. (Right) Overlay of the $^{1}H^{15}N$-TROSY spectra of fully $^{15}N$-labeled HisF in the presence of ProFAR (black) and the $^{1}H^{15}N$-plane of an HNCO spectrum of $^{13}C$,$^{15}N$-labeled HisF, where the Asn are unlabelled, in the presence of ProFAR (red). The signal of F23 is indicated. It is missing from the HNCO spectrum due to the unlabelling of N22. As F23 is the only Phe or Tyr residue that succeeds an Asn residue the lack of this signal in both spectra of the selectively unlabelled HisF variants allows for an unambiguous assignment to F23.

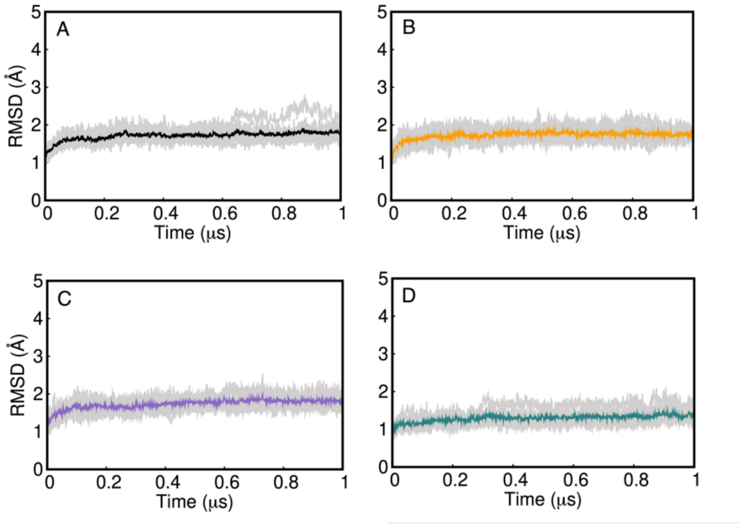


**Figure S14**: **Root mean square deviations (RMSD, Å) of the Cα-atoms during MD simulations of unliganded HisF, initiated from the open conformation of loop1**.
(**A**) wt-HisF, (**B**) HisF-F38A, (**C**) HisF-F23A and (**D**) HisF-G20P. Data was collected every 10 ps from 5 replicas of 1 μs length each. The grey lines show the 5 individual runs, whilst the color solid line shows a rolling average RMSD from all 5 replicas for each system.

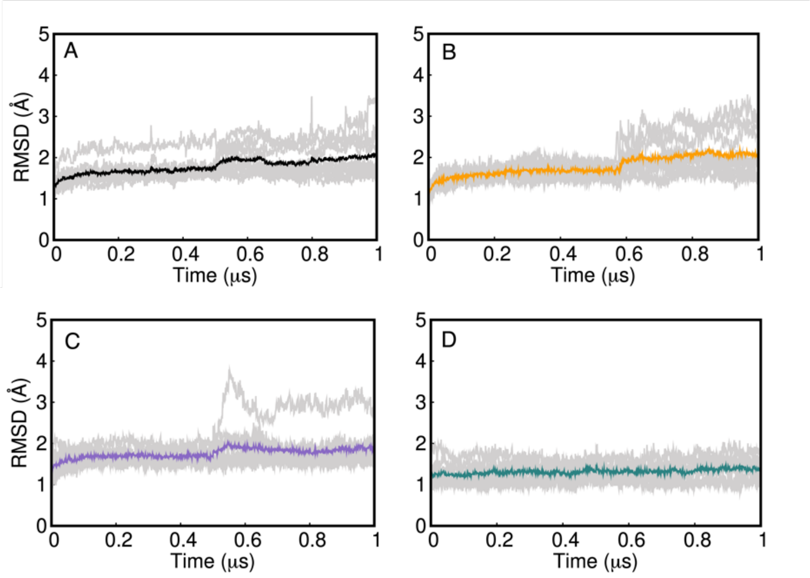


**Figure S15**: **Root mean square deviations (RMSD, Å) of the Cα-atoms during MD simulations of PrFAR-bound HisF, initiated from the open conformation of loop1**.
(**A**) wt-HisF, (**B**) HisF-F38A, (**C**) HisF-F23A and (**D**) HisF-G20P. Data was collected every 10 ps from 5 replicas of 1 μs length each. The grey lines show the 5 individual runs, whilst the color solid line shows a rolling average RMSD from all 5 replicas for each system.

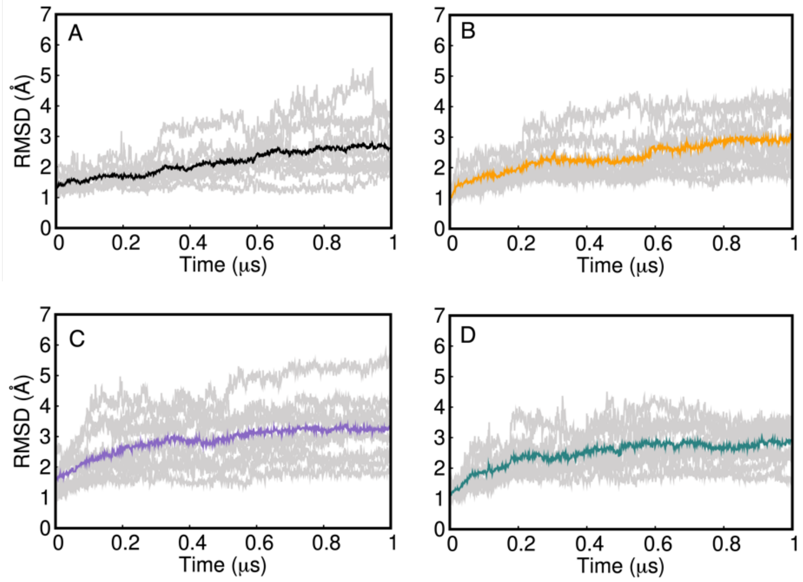


**Figure S16**: **Root mean square deviations (RMSD, Å) of the $C_\alpha$-atoms during MD simulations of PrFAR-bound HisF, initiated from the closed conformation of loop1**.
(**A**) wt-HisF, (**B**) HisF-F38A, (**C**) HisF-F23A and (**D**) HisF-G20P. Data was collected every 10 ps from 5 replicas of 1 μs length each. The grey lines show the 5 individual runs, whilst the color solid line shows a rolling average RMSD from all 5 replicas for each system.

### S2. SUPPLEMENTAL TABLES

##### Table S1: Dissociation constants ($K_D$) determined in fluorescence equilibrium titrations of HisF-CouA.

| | PrFAR | ImGP | | AICAR | |
|---|---|---|---|---|---|
| | | (-AICAR) | (+ AICAR) | (- ImGP) | (+ ImGP) |
| **wt** | 2.4 ± 0.2 µM | 122 ± 17 µM | 70 ± 4.3 µM | 0.97 ± 0.12 mM | 0.45 ± 0.02 mM |
| **F38A** | 2.0 ± 0.5 µM | 94 ± 11 µM | 49 ± 2.0 µM | 1.32 ± 0.15 mM | 0.33 ± 0.01 mM |
| **F23A** | 4.3 ± 0.8 µM | 149 ± 14 µM | 87 ± 17 µM | 1.32 ± 0.20 mM | 0.61 ± 0.08 mM |
| **G20P** | 5.5 ± 0.4 µM | 127 ± 11 µM | 102 ± 10 µM | 1.90 ± 0.87 mM | 0.86 ± 0.11 mM |

$K_D$ values ± SE were determined by fitting the mean ± SEM for at least two technical replicates with equation 3.

##### Table S2: Fluorescence amplitudes (ΔF) determined in fluorescence equilibrium titrations of HisF-CouA and their difference between binary and ternary complexes.

| ImGP | | | AICAR | | |
|---|---|---|---|---|---|
| | $\Delta F_{max}$ (FU) | Diff (+/- AICAR) | | $\Delta F_{max}$ (FU) | Diff (+/- ImGP) |
| **wt** | 163 ± 10 | + 42.9 % | **wt** | 125 ± 8 | + 34.4 % |
| **wt*AICAR** | 233 ± 6 | | **wt*ImGP** | 168 ± 3 | |
| **F38A** | 206 ± 11 | + 50.0 % | **F38A** | 194 ± 13 | + 23.7 % |
| **F38A*AICAR** | 309 ± 4 | | **F38A*ImGP** | 240 ± 4 | |
| **F23A** | 104 ± 5 | - 31.7 % | **F23A** | 121 ± 9 | - 53.7 % |
| **F23A*AICAR** | 71 ± 6 | | **F23A*ImGP** | 56 ± 3 | |
| **G20P** | 224 ± 8 | - 45.1 % | **G20P** | 249 ± 19 | - 47.0 % |
| **G20P*AICAR** | 123 ± 5 | | **G20P*ImGP** | 132 ± 8 | |

$\Delta F_{max}$ values ± SE were determined by fitting the mean ± SEM for at least two technical replicates with equation 3.

##### Table S3: Rate constants for the binding of PrFAR to HisF-CouA at 25°C derived from analysis of secondary plots.

| | **wt** | | **F38A** | |
|---|---|---|---|---|
| $K_D1$ | 6.6 ± 1.1 µM | $K_D$ = 1.1 ± 0.2 µM[1] | 9.0 ± 2.4 µM | $K_D$ = 1.8 ± 0.3 µM[1] |
| $k_{conf}$ | 37.5 ± 1.3 $s^{-1}$ | | 40.1 ± 2.7 $s^{-1}$ | |
| $k_{-conf}$ | 7.8 ± 1.1 $s^{-1}$ | | 9.8 ± 1.6 $s^{-1}$ | |
| | **F23A** | | **G20P** | |
| $k_1$ | 1.3 ± 0.1 $\mu M^{-1} s^{-1}$ | $K_D$ = 11.6 ± 1.2 µM[2] | 2.5 ± 0.1 $\mu M^{-1}s^{-1}$ | $K_D$ = 13.9 ± 0.7 µM[2] |
| $k_{-1}$ | 15.1 ± 1.1 $s^{-1}$ | | 34.8 ± 1.1 $s^{-1}$ | |

The values ± SE for constants $K_D1$, $k_{conf}$, $k_{-conf}$ were determined by fitting the observed rate constants in stopped-flow measurements (**Figure S7E**) to the hyperbolic equation $k_{obs}1 = k_{-conf} + k_{conf}$ *[PrFAR]/$K_{D1}$ + [PrFAR], the values ± SE for the rate constants $k_1$ and $k_{-1}$ were determined by fitting the observed rate constants (**Figure S7F**) to linear equations $k_{obs} = k_{-1} + k_1$ * [PrFAR].

[1] $K_D$ values for induced fit models were calculated using the equation $K_D = K_D1/(1 + k_{conf}/k_{-conf})$[5], SE for $K_D$ values were calculated according to the Gaussian law of error propagation.

[2] $K_D$ values for two-state binding models were calculated using the equation $K_D = k_{-1}/k_1$, SE for $K_D$ values were calculated according to the Gaussian law of error propagation.

**Table S4. Rate constants for the release of ImGP and AICAR from wt-HisF-CouA and HisF-F38A-CouA derived from global fitting analysis.**

| | **wt** | | **F38A** | |
|---|---|---|---|---|
| $\boldsymbol{k_1}$ | 0.18 ± 0.04 $\mu M^{-1}\ s^{-1}$ | $K_D1$ = 242 ± 54 µM | 0.33 ± 0.05 $\mu M^{-1}\ s^{-1}$ | $K_D1$ = 235 ± 36 µM |
| $\boldsymbol{k_{-1}}$ | 44 ± 1 $s^{-1}$ | | 78 ± 2 $s^{-1}$ | |
| $\boldsymbol{k_2}$ | 0.044 ± 0.016 $\mu M^{-1}\ s^{-1}$ | $K_D2$ = 2.05 ± 0.94 mM | 0.085 ± 0.016 $\mu M^{-1}\ s^{-1}$ | $K_D2$ = 1.05 ± 0.20 mM |
| $\boldsymbol{k_{-2}}$ | 90 ± 25 $s^{-1}$ | | 89 ± 2 $s^{-1}$ | |
| $\boldsymbol{k_{conf}}$ | 42 ± 2 $s^{-1}$ | $K_{conf}$ = 0.79 ± 0.04 | 74 ± 5 $s^{-1}$ | $K_{conf}$ = 0.57 ± 0.04 |
| $\boldsymbol{k_{-conf}}$ | 33 ± 1 $s^{-1}$ | | 42 ± 1 $s^{-1}$ | |

The values ± SE for the rate constants $k_1$, $k_{-1}$, $k_2$, $k_{-2}$, $k_{conf}$ and $k_{-conf}$ were obtained in a global fitting analysis according to the model in **Figure S11C**. $K_D$ values were calculated from the rate constants obtained in the global fitting analysis using the equation $K_Dx = k_{-x}/k_x$, $K_{conf} = k_{-conf}/k_{conf}$, SE for $K_D$ values were calculated according to the Gaussian law of error propagation.

**Table S5: Multiple- and single-turnover rates of wt-HisF, HisF-F38A, HisF-F23A, and HisF-G20P.**

| | **Multiple Turnover** | | **Single Turnover** |
|---|---|---|---|
| | $\boldsymbol{k_{cat}}$ **($s^{-1}$)** | $\boldsymbol{K_M^{PrFAR}}$ **(µM)** | $\boldsymbol{k_{obs}}$ **($s^{-1}$)** |
| **wt** | 2.4 ± 0.1 | 3.1 ± 0.5 | 5.7 ± 0.6 |
| **F38A** | 2.6 ± 0.1 | 3.5 ± 0.3 | 5.7 ± 0.3 |
| **F23A** | 0.0074 ± 0.0004 | 19.4 ± 3.9 | 0.0020 ± 0.0002 |
| **G20P** | 0.0017 ± 0.0003 | 6.2 ± 0.4 | 0.0010 ± 0.0002 |

The stopped-flow measurements from which the turnover rates were derived are shown in **Figure 8** (wt-HisF), **Figures S10** (HisF-F38A), **S11** (HisF-F23A) and **S12** (HisF-G20P).

**Table S6. Data collection and refinement statistics for the structures of HisF-G20P and HisF-F23A.**

| **Protein** | **HisF-G20P** | **HisF-F23A** |
|---|---|---|
| **Wavelength (Å)** | 0.99 | 0.99 |
| **Resolution range (Å)** | 47.46 – 1.31 (1.36 – 1.31) | 35.07 – 1.20 (1.24 – 1.20) |
| **Space group** | $P\ 2_1\ 2_1\ 2_1$ | C 1 2 1 |
| **Unit cell** | 44.3, 58.1, 82.4, 90, 90, 90 | 79.3, 44.2, 63.5, 90, 111.8, 90 |
| **Total reflections** | 609236 (29703) | 405073 (35017) |
| **Unique reflections** | 49373 (3705) | 62429 (5730) |
| **Multiplicity** | 12.3 (8.0) | 6.5 (6.0) |
| **Completeness (%)** | 96.16 (73.49) | 96.59 (89.73) |
| **Mean I/sigma(I)** | 12.86 (0.93) | 37.69 (8.88) |
| **Wilson B-factor** | 19.54 | 12.05 |
| $\boldsymbol{R_{merge}}$ | 0.091 (1.37) | 0.026 (0.18) |
| $\boldsymbol{R_{meas}}$ | 0.095 (1.46) | 0.028 (0.20) |
| $\boldsymbol{R_{pim}}$ | 0.026 (0.50) | 0.011 (0.079) |
| $\boldsymbol{CC_{1/2}}$ | 1.00 (0.37) | 1.00 (0.98) |
| **CC*** | 1 (0.73) | 1 (0.996) |
| **Reflections used in refinement** | 49357 (3704) | 62307 (5730) |
| **Reflections used for $\boldsymbol{R_{free}}$** | 1999 (150) | 2012 (182) |
| $\boldsymbol{R_{work}}$ | 0.20 (0.43) | 0.20 (0.22) |

| | | |
|---|---|---|
| **$R_{free}$** | 0.22 (0.44) | 0.21 (0.21) |
| **$CC_{work}$** | 0.97 (0.66) | 0.95 (0.90) |
| **$CC_{free}$** | 0.95 (0.46) | 0.91 (0.83) |
| **Number of atoms** | 2180 | 2245 |
| **macromolecules** | 1929 | 1954 |
| **ligands** | 0 | 10 |
| **solvent** | 251 | 281 |
| **Protein residues** | 250 | 255 |
| **RMS (bonds)** | 0.006 | 0.005 |
| **RMS (angles)** | 1.15 | 0.79 |
| **Ramachandran favored (%)** | 98.79 | 96.41 |
| **Ramachandran allowed (%)** | 1.21 | 3.59 |
| **Ramachandran outliers (%)** | 0.00 | 0.00 |
| **Rotamer outliers (%)** | 0.48 | 0.00 |
| **Clashscore** | 4.37 | 6.54 |
| **Average B-factor** | 24.08 | 17.96 |
| **macromolecules** | 22.91 | 16.88 |
| **ligands** | - | 14.00 |
| **solvent** | 33.08 | 25.62 |

Statistics for the highest resolution shell are shown in parentheses.

**Table S7. Non-standard force field parameters used to describe the substrate PrFAR in the molecular dynamics simulations.[a]**

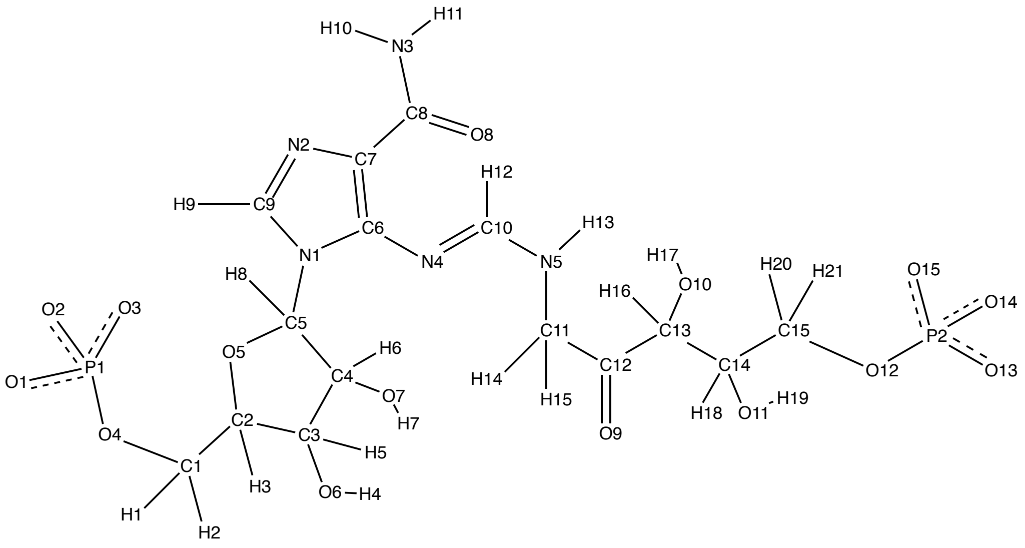


| Atom Name | Atom Type | Charge |
|---|---|---|
| O1 | o | -0.995514 |
| O2 | o | -0.995514 |
| O3 | o | -0.995514 |
| P1 | p5 | 1.548494 |
| O4 | os | -0.554789 |
| C1 | c3 | 0.146878 |
| H1 | h1 | -0.015493 |
| H2 | h1 | -0.015493 |
| C2 | c3 | 0.100325 |
| H3 | h1 | 0.074478 |
| C3 | c3 | 0.174417 |
| O6 | oh | -0.707284 |
| H4 | ho | 0.401459 |
| H5 | h1 | 0.03885 |
| C4 | c3 | 0.346713 |
| H6 | h1 | 0.021574 |
| O7 | oh | -0.726354 |
| H7 | ho | 0.432551 |
| C5 | c3 | 0.233948 |

| Atom Name | Atom Type | Charge |
|---|---|---|
| N2 | nd | -0.498464 |
| C9 | cc | 0.003374 |
| H9 | h5 | 0.320119 |
| N4 | ne | -0.491428 |
| C10 | c2 | 0.346756 |
| H12 | h5 | 0.050122 |
| N5 | nh | -0.330124 |
| H13 | hn | 0.283768 |
| C11 | c3 | -0.322921 |
| H14 | h1 | 0.108211 |
| H15 | h1 | 0.108211 |
| C12 | c | 0.796782 |
| O9 | o | -0.609654 |
| C13 | c3 | 0.132262 |
| H16 | h1 | 0.020844 |
| O10 | oh | -0.841934 |
| H17 | ho | 0.588213 |
| C14 | c3 | 0.265635 |
| H18 | h1 | -0.005259 |

| | | | | | |
|---|---|---|---|---|---|
| H8 | h2 | 0.043035 | O11 | oh | -0.724352 |
| O5 | os | -0.480638 | H19 | ho | 0.423614 |
| N1 | na | 0.087669 | C15 | c3 | 0.156364 |
| C6 | cc | 0.139799 | H20 | h1 | -0.002368 |
| C7 | cd | -0.030756 | H21 | h1 | -0.002368 |
| C8 | c | 0.754656 | O12 | os | -0.585436 |
| N3 | n | -0.960604 | P2 | p5 | 1.633481 |
| H11 | hn | 0.377226 | O13 | o | -1.003258 |
| H10 | hn | 0.377226 | O14 | o | -1.003258 |
| O8 | o | -0.635019 | O15 | o | -1.003258 |

[a] All parameters were obtained using the General AMBER Force Field 2 (GAFF2) as described in the **Supplemental Methods** section.

**Table S8. Distance restraints employed during all MD simulations to retain the PrFAR substrate in the binding pocket.[a]**

| Distance Restraints | | | | | | | |
|---|---|---|---|---|---|---|---|
| Atom 1 | Atom 2 | Distances (Å) | | | | Restraints (kcal mol$^{-1}$ A$^{-2}$) | |
| | | r[1] | r[2] | r[3] | r[4] | r*k*[2] | r*k*[3] |
| PRFAR@O2 | G82@N | 2.0 | 3.0 | 5.0 | 10.0 | 0.0 | 10.0 |
| PRFAR@O3 | T104@N | 2.0 | 3.0 | 5.0 | 10.0 | 0.0 | 10.0 |
| PRFAR@C4 | S101@OG | 2.0 | 3.0 | 5.0 | 10.0 | 0.0 | 10.0 |
| PRFAR@O9 | D11@OD1 | 2.0 | 3.0 | 5.0 | 10.0 | 0.0 | 10.0 |
| PRFAR@O11 | S201@OG | 2.0 | 3.0 | 5.0 | 10.0 | 0.0 | 10.0 |
| PRFAR@O14 | S225@H | 2.0 | 3.0 | 5.0 | 10.0 | 0.0 | 10.0 |
| PRFAR@O15 | G203@N | 2.0 | 3.0 | 5.0 | 10.0 | 0.0 | 10.0 |

[a] $r^1$, $r^2$, $r^3$ and $r^4$ values indicate the four positions used to define the distance restraints between the two atoms. $rk^2$ and $rk^3$ define the force constants for the left and right parabolas, respectively. Note that distance restraints are wall restraints that kick when the two atoms are 5.0 Å or greater apart. No restraints were placed between the PRFAR substrate and any loop1 residue.

**Table S9. Distance restraints employed during all MD simulations to avoid interactions between neutralizing counterions and the PrFAR substrate within the active site.[a]**

| Distance Restraints | | | | | | | |
|---|---|---|---|---|---|---|---|
| Atom 1 | Atom 2 | Distances (Å) | | | | Restraints (kcal mol$^{-1}$ A$^{-2}$) | |
| | | r[1] | r[2] | r[3] | r[4] | r*k*[2] | r*k*[3] |
| Na+255 | D98@OD1 | 2.0 | 3.0 | 6.0 | 7.5 | 5.0 | 5.0 |
| Na+256 | D31@OD2 | 2.0 | 3.0 | 6.0 | 7.5 | 5.0 | 5.0 |
| Na+257 | E71@OE1 | 2.0 | 3.0 | 6.0 | 7.5 | 5.0 | 5.0 |
| Na+258 | E236@OE1 | 2.0 | 3.0 | 6.0 | 7.5 | 5.0 | 5.0 |
| Na+259 | D74@OD1 | 2.0 | 3.0 | 6.0 | 7.5 | 5.0 | 5.0 |
| Na+260 | E137@OE1 | 2.0 | 3.0 | 6.0 | 7.5 | 5.0 | 5.0 |
| Na+261 | E161@OE2 | 2.0 | 3.0 | 6.0 | 7.5 | 5.0 | 5.0 |
| Na+262 | E91@OE1 | 2.0 | 3.0 | 6.0 | 7.5 | 5.0 | 5.0 |
| Na+263 | E239@OE1 | 2.0 | 3.0 | 6.0 | 7.5 | 5.0 | 5.0 |
| Na+264 | D135@OD1 | 2.0 | 3.0 | 6.0 | 7.5 | 5.0 | 5.0 |
| Na+265 | E41@OE1 | 2.0 | 3.0 | 6.0 | 7.5 | 5.0 | 5.0 |
| Na+266 | E57@OE1 | 2.0 | 3.0 | 6.0 | 7.5 | 5.0 | 5.0 |

[a] $r^1$, $r^2$, $r^3$ and $r^4$ values indicate the four positions used to define the distance restraints between the two atoms. $rk^2$ and $rk^3$ define the force constants for the left and right parabolas, respectively. Note that distance restraints are harmonic restraints that kick when the two atoms are closer than 2.0 Å or greater than 6.0 Å.

### S3. SUPPLEMENTAL METHODS

#### Script file DynaFit, binding of AICAR/ImGP to wt-HisF CouA

```
[task]
  data = progress
  task = fit
  model = induced fit1

[mechanism]
  E + I <==> EI   :   k1  k-1
  E + A <==> EA   :   k2  k-2
  EA + I <==> EAI :   k1  k-1
  EI + A <==> EAI :   k2  k-2
  EAI <==> EAI*   :   kconf k-conf

[constants]                          ; µM, s
  k1 = 0.8?
  k-1 = 86?
  k2 = 0.2?
  k-2 = 90?
  kconf = 150?
  k-conf = 35?

[responses]
  E = 13.2
  I = 0.01
  EI = 12.7?
  EA = 13.0?
  EAI = 6.4
  EAI* = 4.6

[data]
  directory $IN_DIR
  extension csv
  delay 0.002

file AICAR0 | conc E = 0.05 | offset 0.97 ?
file AICAR1 | conc E = 0.05, A = 250 | offset 0.97 ?
file AICAR2 | conc E = 0.05, A = 1000 | offset 0.97 ?
file ImGP0 | conc E = 0.05 | offset 0.97 ?
file ImGP1 | conc E = 0.05, I = 100 | offset 0.97 ?
file ImGP2 | conc E = 0.05, I = 250 | offset 0.97 ?
file Mix1  | conc E = 0.05, A = 250, I = 100 | offset 0.97?
file Mix2  | conc E = 0.05, A = 250, I = 250 | offset 0.97 ?
file Mix3  | conc E = 0.05, A = 1000, I = 100 | offset 0.97 ?
file Mix4  | conc E = 0.05, A = 1000, I = 250 | offset 0.97 ?

file AIC_ImGP1 | equilibrate E = 0.1, A = 2500, dilute 0.5 | conc I = 0 | offset 0.97 ?
file AIC_ImGP2 | equilibrate E = 0.1, A = 2500, dilute 0.5 | conc I = 100 | offset 0.97 ?
file AIC_ImGP3 | equilibrate E = 0.1, A = 2500, dilute 0.5 | conc I = 250 | offset 0.97 ?
file Im_AICAR1 | equilibrate E = 0.1, I = 400, dilute 0.5 | conc A  = 0 | offset 0.97 ?
file Im_AICAR2 | equilibrate E = 0.1, I = 400, dilute 0.5 | conc A  = 250 | offset 0.97 ?
file Im_AICAR3 | equilibrate E = 0.1, I = 400, dilute 0.5 | conc A = 1000 | offset 0.97 ?

[output]
  directory $OUT_DIR
[end]
```

#### Additional Simulation Details

*System Preparation for Conventional Molecular Dynamics Simulations*

Structure preparation for molecular dynamics simulations was performed using AmberTools22[6], and all simulations were performed using the AMBER ff14SB force field[7]. All crystallographic water molecules and ions were stripped from the crystal structure, and the protein was resolvated in a truncated octahedral water box of TIP3P[8] water molecules, with solvent extending 10 Å from the protein in all directions. The most likely protonation state of ionizable side chains in the system was determined using AmberTools22 and cross checked using PROPKA3.5.1[9]. Based on this, all residues were kept in their expected standard protonation states at physiological pH. For each system, sodium counterions were added to the simulation to ensure overall charge neutrality of the system. The partial charges of the ligand PrFAR were computed based on Restrained Electrostatic Potential[10] (RESP) fitting using Antechamber[11], with the electrostatic potential of the substrate calculated at the M062X/6-31+G(d) level of theory, in diethylether (decribed by the conductor-like polarizable continuum model[12] (CPCM)). The electrostatic potential was calculated after performing an initial geometry optimization of the structure at the same level of theory, using Gaussian 16 Rev. C.01[13]. All other parameters to describe the substrate PrFAR were obtained using the General AMBER Force Field 2[14] (GAFF2). All non-standard parameters to describe the substrate are provided in **Table S7** and in the Zenodo data package available at DOI: 10.5281/zenodo.1211377.

Finally, in the case of simulations of HisF variants in complex with PrFAR, positional restraints were (1) applied between the ligand and the active site residues to ensure that the ligand remains in the binding site, and (2) the neutralizing counterions in order to keep them away from the active site, otherwise they showed a tendency to try to bind to the phosphodianion groups of the substrate and unphysically disrupt loop dynamics. A summary of these restraints is provided in **Tables S8** and **S9**.

*Molecular Dynamics Equilibration Procedure*

Following system set-up, ten replicas of each system were generated by assigning different random velocities for each system following a standard Boltzmann distribution. All equilibration simulations were performed using a 1 fs timestep, and applying the SHAKE[15] algorithm to constrain all bonds involving a hydrogen atom. The systems were then equilibrated using the following steps: (1) 1000 steps of steepest descent energy minimization at 0K to remove initial bad contacts in the starting structure. (2) Gradual heating of the system from 100 to 300K over 1 ns of equilibration time. During this heating step, all protein and PrFAR (if bound) atoms were restrained using 100 kcal $mol^{-1}$ $Å^{-2}$ harmonic restraints. (3) A 1 ns long NPT simulation, keeping the restraints from step 2. (4) 1 ns long NPT simulation, reducing the restraints on the protein and PrFAR atoms to 10 kcal $mol^{-1}$ $Å^{-2}$. (5) A further 1 ns long NPT simulation, with a 10 kcal $mol^{-1}$ $Å^{-2}$ restrained applied only to the heavy atoms of the protein and PRFAR. (6) A further 1 ns long NPT simulation with the restraint on the protein/PrFAR heavy atoms reduced to 1 kcal $mol^{-1}$ $Å^{-2}$. (7) A final 1 ns long restrained NPT simulation with the protein/PrFAR heavy atom restraint reduced to 0.1 kcal $mol^{-1}$ $Å^{-2}$. (8) A final 1 ns unrestrained NPT simulation as the last step of the equilibration before production simulations. All NVT simulations used Langevin temperature control (collision frequency of 1 $ps^{-1}$), while all NPT simulations used both Langevin temperature control (collision frequency of 1 $ps^{-1}$) and a Berendsen barostat (1 ps pressure relaxation time). Equilibration of these trajectories is shown in **Figures S15-S17**.

*Simulation Analysis*

All MD analyses in this work were performed using the CPPTRAJ[16] module of AmberTools23[6], unless otherwise indicated. Simulation snapshots were extracted for analysis every 1ns of the trajectory, and the analyses presented in this work are average values and standard deviations over 10 x 1 μs trajectories per system.

The mobility of loop1 (residues 15-30) was analyzed by computing the first 5 normal modes of the system, by calculating the mass-weighted coordinate covariance matrix followed by the root mean square fluctuation (RMSF) of the $C_\alpha$-atoms.

2D histograms of loop1 motion (**Figures 5**) were generated as a function of the RMSD of loop1 to its closed conformation in the HisF/HisH crystal structure (PDB ID: 7ac8[40], chains E and F) and the distance RMSD (dRMSD) of key non-covalent interactions between loop1 and the protein scaffold, projected onto a single vector. The interactions were identified using standard interaction analysis as implemented in the Key Interaction Networks (KIN)[3] package, and were visually confirmed using PyMOL.[17]

Based on this analysis, the following interactions were identified in the open crystal structure of the wt-HisF (PDB ID: 1thf) and were used to define the dRMSD: Phe23-Phe38, Phe23-Tyr39, Phe23-Arg230, Asn25-Arg230, Arg27-Glu34, Asp28-Glu34, Asp31-Val33, Asp31-Glu34 and Asp31-Leu35. We specifically selected interactions between the protein and the open conformation (rather than the closed conformation) of loop1 for our dRMSD calculations, in order to be able to effectively distinguish between "open" conformations of the loop which maintain interactions with the protein scaffold, and detached conformations of the loop which lose these interactions, as described in **Figure 1**.

Non-covalent interactions between the loop1 and the substrate PrFAR were analyzed for representative structures using NCIWeb[18]. Finally, PyMOL[17] was used for visualization analysis.

13. Frisch, M. J.; Trucks, G. W.; Schlegel, H. B.; Scuseria, G. E.; Robb, M. A.; Cheeseman, J. R.; Scalmani, G.; Barone, V.; Petersson, G. A.; Nakatsuji, H.; Li, X.; Caricato, M.; Marenich, A. V.; Bloino, J.; Janesko, B. G.; Gomperts, R.; Mennucci, B.; Hratchian, H. P.; Ortiz, J. V.; Izmaylov, A. F.; Sonnenberg, J. L.; Williams; Ding, F.; Lipparini, F.; Egidi, F.; Goings, J.; Peng, B.; Petrone, A.; Henderson, T.; Ranasinghe, D.; Zakrzewski, V. G.; Gao, J.; Rega, N.; Zheng, G.; Liang, W.; Hada, M.; Ehara, M.; Toyota, K.; Fukuda, R.; Hasegawa, J.; Ishida, M.; Nakajima, T.; Honda, Y.; Kitao, O.; Nakai, H.; Vreven, T.; Throssell, K.; Montgomery Jr., J. A.; Peralta, J. E.; Ogliaro, F.; Bearpark, M. J.; Heyd, J. J.; Brothers, E. N.; Kudin, K. N.; Staroverov, V. N.; Keith, T. A.; Kobayashi, R.; Normand, J.; Raghavachari, K.; Rendell, A. P.; Burant, J. C.;

Iyengar, S. S.; Tomasi, J.; Cossi, M.; Millam, J. M.; Klene, M.; Adamo, C.; Cammi, R.; Ochterski, J. W.; Martin, R. L.; Morokuma, K.; Farkas, O.; Foresman, J. B.; Fox, D. J. *Gaussian 16 Rev. C.01*, Wallingford, CT, 2016.
14. Wang, J. M.; Wolf, R. M.; Caldwell, J. W.; Kollman, P. A.; Case, D. A., Development and Testing of a General Amber Force Field. *J. Comput. Chem.* **2004,** *25*, 1157-1174.
15. Ryckaert, J.-P.; Ciccotti, G.; Berendsen, H. J. C., Numerical Integration of the Cartesian Equations of Motion of a System With Constraints: Molecular Dynamics of *n*-Alkanes. *J. Comp. Physics* **1977,** *23*, 327-341.
16. Roe, D. R.; Cheatham, T. E., 3rd, PTRAJ and CPPTRAJ: Software for Processing and Analysis of Molecular Dynamics Trajectory Data. *J Chem Theory Comput* **2013,** *9*, 3084-3095.
17. Schrodinger, LLC, The PyMOL Molecular Graphics System, Version 1.8. 2015.
18. Novoa, T.; Laplaza, R.; Peccati, F.; Fuster, F.; Contreras-Garcia, J., The NCIWEB Server: A Novel Implementation of the Noncovalent Interactions Index for Biomolecular Systems. *J. Chem. Inf. Model.* **2023,** *63*, 4483-4489.
